# The endophilin curvature-sensitive motif requires electrostatic guidance to recycle synaptic vesicles *in vivo*

**DOI:** 10.1101/2021.10.20.465157

**Authors:** Lin Zhang, Yu Wang, Yongming Dong, Aaradhya Pant, Yan Liu, Laura Masserman, Richard N McLaughlin, Jihong Bai

## Abstract

Curvature-sensing mechanisms assist proteins in executing particular actions on various membrane organelles. Here, we investigated the functional specificity of curvature-sensing amphipathic motifs through the study of endophilin, an endocytic protein for synaptic vesicle recycling. We generated chimeric endophilin proteins by replacing the endophilin amphipathic motif H0 with other curvature-sensing amphipathic motifs. We found that the role of amphipathic motifs cannot simply be extrapolated from the identity of their parental proteins. For example, the amphipathic motif of the nuclear pore complex protein NUP133 functionally replaced the synaptic role of endophilin H0. Interestingly, non-functional endophilin chimeras had similar defects – producing fewer synaptic vesicles but more endosomes – indicating that the curvature-sensing motifs in these chimeras have a common deficiency at reforming synaptic vesicles. Finally, we converted non-functional endophilin chimeras into functional proteins by changing the cationic property of amphipathic motifs, setting a precedent for reprogramming the functional specificity of curvature-sensing motifs *in vivo*.

## Introduction

Eukaryotic cells manage a network of membrane organelles that perform specific functions. Research on curvature-sensing mechanisms has intensified in recent years, as these principles show promise in coupling membrane morphology to biological activities (Antonny, 2011; Jarsch et al., 2016; McMahon and Gallop, 2005; Prinz and Hinshaw, 2009; Shibata et al., 2009). Such efforts have led to the identification of many curvature-sensing domains and motifs in proteins with various cellular functions, raising a possibility that curvature-sensing mechanisms assist proteins to perform specific activities on distinct membrane organelles. Yet, how exactly these mechanisms provide specific guidance to proteins *in vivo* remains largely unknown.

One class of curvature-sensing modules consists of amphipathic motifs, which are short membrane-binding sequences found in many proteins and peptides (Bigay and Antonny, 2012; Drin and Antonny, 2010; Gimenez-Andres et al., 2018; Jarsch *et al*., 2016; Shibata *et al*., 2009). In solution, the amphipathic motifs are disordered. Upon contact with curved membranes, they fold into a helical structure, with polar and nonpolar residues segregated into two faces of the helix (Cornell and Taneva, 2006; Kaiser and Kezdy, 1983; 1984; Sargent et al., 1988). The amphipathic helices lay parallel onto membranes with residues in the nonpolar face embedded within the membrane hydrophobic core, and residues in the polar face oriented towards the aqueous cytoplasm and lipid headgroups (Hristova et al., 1999). The hydrophobic residues in the nonpolar face recognize lipid packing defects of curved membranes, consequently enabling curvature-dependent binding of amphipathic motifs to membranes (Bartels et al., 2010; Cui et al., 2011; Gonzalez-Rubio et al., 2011; Hatzakis et al., 2009; Vanni et al., 2013).

While attempting to understand the diversity among amphipathic motifs, biochemical studies were conducted which found a number of factors that influence the curvature sensitivity of these motifs *in vitro*. For instance, the composition of amino acid residues in the polar face has a strong impact on curvature sensitivity. The Amphipathic-Lipid-Packing-Sensor (ALPS) motifs, which have polar faces that mainly contain serine and threonine residues, are highly sensitive to membrane curvature (Drin and Antonny, 2010; Drin et al., 2007). In comparison, amphipathic motifs that have polar faces with abundant basic residues, and can thus bind anionic membranes through electrostatic interactions, are therefore thought to be less selective to curvature (Drin *et al*., 2007). Besides hydrophilic and charged residues in the polar face, the curvature-dependent membrane interactions of amphipathic motifs could be tuned by a number of additional parameters including the size and the density of hydrophobic residues in the nonpolar face (Bigay et al., 2005; Boucrot et al., 2012; Farsad et al., 2001; Ford et al., 2002; Gallop et al., 2006; Krauss et al., 2008; Lee et al., 2005; Masuda et al., 2006; Suresh and Edwardson, 2010), the length of the amphipathic motifs (Braun et al., 2017; Copic et al., 2018; McLean et al., 1991), the surrounding protein backbones (Doucet et al., 2015), and posttranslational modifications (Karanasios et al., 2010). These findings show a high degree of biochemical distinctions among amphipathic motifs, suggesting that these motifs could be further sorted into subgroups *in vitro* (Gimenez-Andres *et al*., 2018). However, little is known about the level of functional specificity these motifs display *in vivo*. For example, can an amphipathic motif be replaced by another one without leading to functional deficiency? Do amphipathic motifs with different *in vitro* properties always show clear functional distinction *in vivo*? What key features need to be altered to make a non-functional amphipathic motif into a functional one? These are major questions to be answered.

Endophilin-A (hereafter endophilin) is one of the best characterized endocytic proteins in neurons (Bai et al., 2010; Dickman et al., 2005; Kittelmann et al., 2013; Kroll et al., 2019; Milosevic et al., 2011; Schuske et al., 2003; Verstreken et al., 2002; Watanabe et al., 2018) and non-neuronal cells (Boucrot et al., 2015; Ferreira and Boucrot, 2018; Poudel et al., 2018; Renard et al., 2015; Renard et al., 2020). It harbors an amphipathic motif H0 at its N-terminus, a Bin-Amphiphysin-Rvs (BAR) domain in the middle, and an SH3 domain at its C-terminus (Farsad *et al*., 2001; Gallop *et al*., 2006; Peter et al., 2004; Ringstad et al., 1997; Weissenhorn, 2005). Removal of endophilin from neurons leads to profound synaptic defects including a reduced pool of synaptic vesicles, decreased levels of synaptic transmission, and slowed recycling of synaptic vesicle proteins and membranes (Bai *et al*., 2010; Dickman *et al*., 2005; Kittelmann *et al*., 2013; Kroll *et al*., 2019; Milosevic *et al*., 2011; Schuske *et al*., 2003; Verstreken *et al*., 2002; Watanabe *et al*., 2018). The importance of endophilin H0 for supporting endocytosis is well established. Truncated endophilin that lacks the H0 motif folds correctly (Cui et al., 2013; Poudel et al., 2016), but it fails to sustain synaptic vesicle recycling in neurons (Bai *et al*., 2010), and it loses the ability to promote endocytic uptake of membranes in non-neuronal cells (Boucrot *et al*., 2015). Like other amphipathic motifs, endophilin H0 is disordered in solution (Bhatia et al., 2009; Gallop *et al*., 2006; Low et al., 2008). It folds rapidly into an amphipathic helix on membranes containing anionic lipids (Capraro et al., 2013; Poudel *et al*., 2016), with side chains of nonpolar residues penetrating into the lipid bilayer (Cui *et al*., 2011; Jao et al., 2010). Because of the functional importance of the H0 motif, extensive effort has been put forward to elucidate the biochemical and biophysical basis for the interactions between endophilin H0 and membranes (Ambroso et al., 2014; Bhatia *et al*., 2009; Blood et al., 2008; Boucrot *et al*., 2012; Capraro *et al*., 2013; Chen et al., 2016b; Cui *et al*., 2011; Cui *et al*., 2013; Hatzakis *et al*., 2009; Low *et al*., 2008; Poudel *et al*., 2016). It is generally agreed that the H0 motif possesses the curvature-sensing activity essential for endophilin to bind curved membranes. The insertion of amphipathic H0 assists endophilin to generate membrane curvature by actively bending membranes (Campelo et al., 2008; Low *et al*., 2008; Poudel *et al*., 2016; Sodt and Pastor, 2014), passively stabilizing lipid packing defects presented on bent membranes (Bhatia *et al*., 2009; Chen *et al*., 2016b; Cui *et al*., 2011; Hatzakis *et al*., 2009; Poudel *et al*., 2016), or enhancing steric repulsion arising from protein crowding on membranes (Chen et al., 2016a; Stachowiak et al., 2012). In addition, the activity of endophilin H0 can be tuned by phosphorylation (Kaneko et al., 2005) and its interaction with the SH3 domain of endophilin (Chen et al., 2014; Vazquez et al., 2013; Wang et al., 2008), which are modulatory mechanisms requiring specific protein sequences. Collectively, these studies suggest that the H0 motif is a special amphipathic motif necessary for the activity of endophilin *in vitro*. However, the specificity of endophilin H0 has not been examined *in vivo*.

In this study, we generated endophilin chimeras by replacing endophilin H0 with other curvature-sensing amphipathic helical motifs. Our results showed that only a subset of amphipathic motifs could support endophilin’s synaptic activity in neurons, demonstrating functional specificity among amphipathic motifs *in vivo*. While endophilin H0 could be functionally replaced by an ALPS motif of the nuclear pore complex protein NUP133, another ALPS motif, from the oxysterol-binding protein KES1, failed to substitute for endophilin H0. Therefore, the specificity of an amphipathic motif cannot be simply extrapolated from the function of its parental protein. Through ultrastructural analyses, we found that endophilin chimeras carrying non-functional curvature-sensing motifs induce the abnormal accumulation of endosome-like structures at synapses, suggesting a common defect in the process of resolving endosomes into synaptic vesicles. Finally, we successfully converted non-functional endophilin chimeras into functional ones by changing the cationic property of the amphipathic motifs. We showed that the functional improvement was due to changes of the net charge levels instead of the exact position of charges, the type of mutations, or the amino acid sequences of the amphipathic motifs. Collectively, this study demonstrated for the first time how electrostatic and curvature-sensing mechanisms of H0 work together to provide specific guidance to endophilin function *in vivo*.

## Results

### The linker region between rEndoA1 H0 and the BAR domain tolerates changes

In support of a conserved function of endophilin in invertebrate and vertebrate neurons, we found that neuronal expression of the GFP-tagged full-length rat endophilin A1 (rEndoA1, Fig.1A), using a single-copy transgene (*snb-1p::rEndoA1::gfp*), fully restored the locomotion rates (Fig.1B) and synaptic currents at neuromuscular junctions in *unc-57 endophilin null* mutant worms (Fig.1C, 1D). On the other hand, removal of the H0 motif from rEndoA1 (ΔH0) completely abolished the rescue activity of rEndoA1 in *unc-57* mutants (Fig.S1).

**Figure 1.**
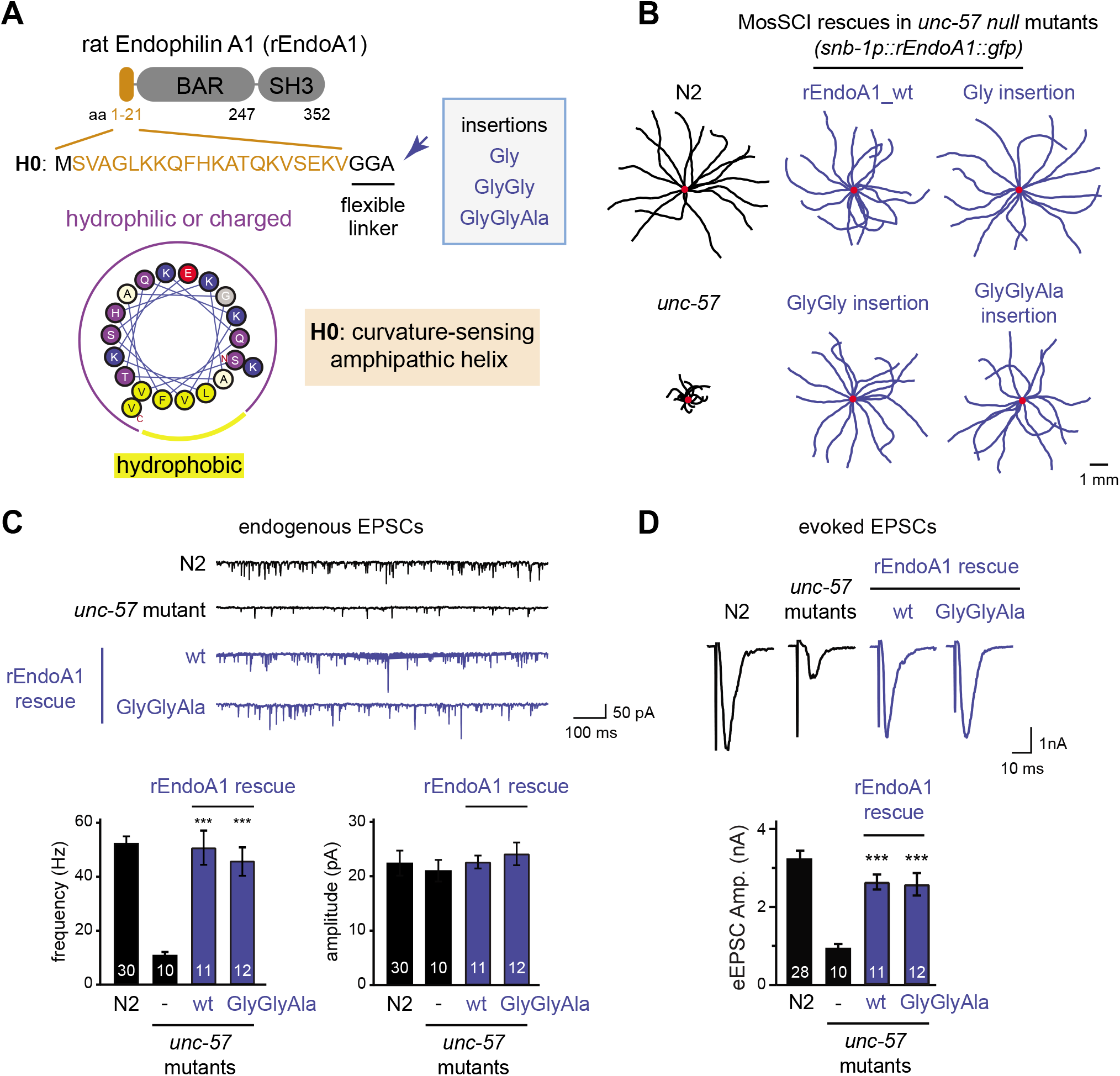
The linker between the N-terminal amphipathic helix H0 and the BAR domain of Endophilin tolerates changes. **(A)** *Upper panel* - Schematic diagram showing the domain structure of rat Endophilin A1 (rEndoA1). The amino acid sequence of the N-terminal amphipathic helix H0 (residues 1-21) is shown in brown, followed by an endogenous flexible linker (GGA) that connects endophilin H0 to the BAR domain. The blue arrow indicates insertions of amino acid residues after the endogenous linker. *Lower panel* – A helical wheel presentation of the rEndoA1_H0 motif. Amino acid residues are colored as following; yellow, hydrophobic; blue, basic; red, acidic; purple, serine, threonine, asparagine, glutamine and histidine; grey, glycine; light yellow, alanine. **(B-D)** Mutant versions of rEndoA1 that carry insertions between endophilin H0 and the BAR domain were assayed for *in vivo* function. A pan-neuronal promoter *snb-1p* was used to drive transgene expression. Single-copy transgenes *(snb-1p::rEndoA1::gfp)* encoding GFP-tagged rEndoA1 variants were introduced into *endophilin unc-57(e406)* mutant worms. Representative trajectories (15 animals) of 30s locomotion for each genotype are shown in **(B)**. The starting points for each trajectory were aligned (indicated by red dots) for clarity. The scale bar indicates 1 mm. Electrophysiological recordings were used to determine endogenous and evoked excitatory postsynaptic currents (EPSCs) at neuromuscular junctions. Representative traces and summary data for endogenous EPSC rates and amplitudes were shown in **(C)**, and evoked EPSC traces and summary data were shown in **(D)**, for indicated genotypes. ***, p < 0.001; when compared to *endophilin unc-57 mutants*. The number of worms analyzed for each genotype is indicated in the bar graphs. Error bars represent standard error of the mean (SEM).

To understand how the N-terminal amphipathic helix H0 works in the context of full-length endophilin, we examined the linker region between H0 and the BAR domain. In wild type rEndoA1, the H0 motif is connected to the BAR domain of endophilin via a short GlyGlyAla linker (Fig.1A), suggesting a flexible nature of the connection. To determine whether the linker requires a particular length of amino acid residues, we generated mutant versions of rEndoA1 that carry additional residues of Gly, GlyGly, and GlyGlyAla before the BAR domain (indicated by the arrow in Fig.1A). Our data showed that mutant rEndoA1 with altered linkers exhibited similar levels of rescue activity in restoring locomotion rates (Fig.1B), the frequency of endogenous excitatory postsynaptic currents (EPSCs) (Fig.1C), and the amplitude of evoked EPSCs in response to electrical stimulation (Fig.1D). Together, these data indicate that the flexible linker that connects the amphipathic helix H0 to the rEndoA1 BAR domain can tolerate changes to its amino acid residue sequence.

### Endophilin H0 carries functional specificity in vivo

We next sought to examine the functional specificity of endophilin H0 *in vivo*. To this end, we generated an array of rEndoA1 chimeras that carry amphipathic motifs from various proteins in place of endophilin H0 (Fig. 2A-B). The amphipathic motifs grafted into these endophilin chimeras have different primary sequences. However, like endophilin H0, they are all known curvature sensors as shown by biochemical studies, *i.e.*, they preferentially bind highly curved membranes *in vitro* (Drin and Antonny, 2010). The rEndoA1 chimeras were expressed in neurons using single-copy transgenes, and the expression levels of rEndoA1 chimeras were measured by determining the GFP fluorescence intensity around the *C. elegans* nerve ring (a tight bundle of neuronal processes of sensory and interneurons). Imaging results showed that rEndoA1 chimeras were expressed in neurons at similar levels to wild type rEndoA1 (Fig.S2).

**Figure 2.**
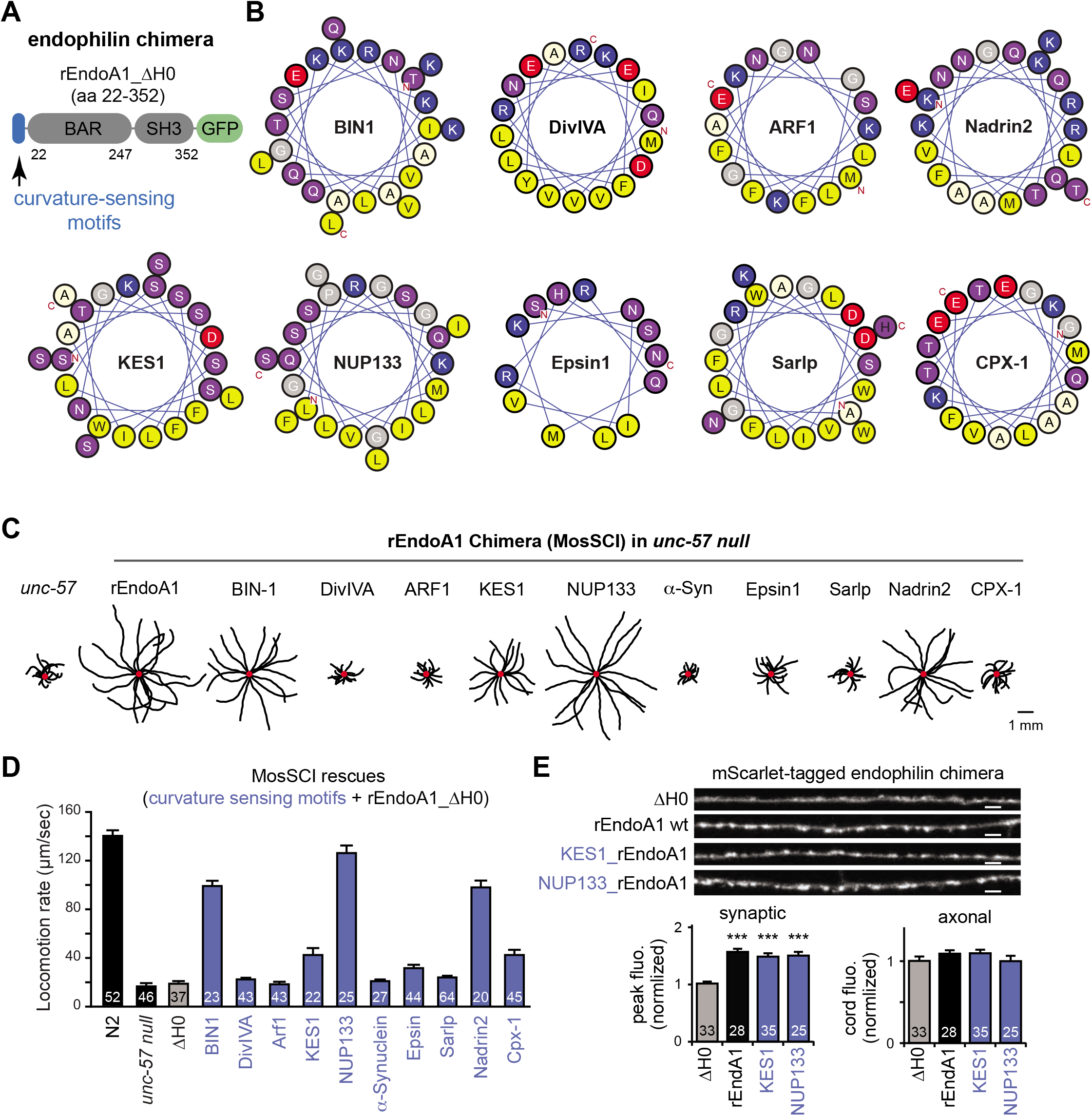
Endophilin chimera carrying amphipathic helices from various proteins displayed functional specificity *in vivo*. **(A)** Schematic diagram showing the domain structure of chimeric versions of rEndoA1 (tagged with GFP at C-terminus). The N-terminal H0 motif of rEndoA1 (colored in blue) was replaced by curvature-sensing amphipathic helices from various proteins. Helical wheel presentations of amphipathic helices used for generating endophilin chimera were shown in **(B)**. Amino acid residues were colored as described in the legend of Figure 1. Single-copy transgenes *(snb-1p::rEndoA1_chimera::gfp)* were introduced into *endophilin unc-57(e406)* mutant worms to drive expression of GFP-tagged rEndoA1 chimera in neurons. **(C)** Representative trajectories (15 animals) of 30s locomotion were shown for indicated genotype. The starting points for each trajectory were aligned (indicated by red dots) for clarity. The scale bar indicates 1 mm. **(D)** Summary data for locomotion rates were plotted for indicated genotypes. **(E)** Fluorescence microscopy was used to examine the distribution of rEndoA1 proteins at synapses and in axons. Representative images and summary data for wild type rEndoA1, mutant rEndoA1 lacking the H0 motif (ΔH0), and chimeric rEndoA1 proteins carrying amphipathic curvature-sensing motifs from NUP133 and KES1 were shown. ***, p < 0.001; when compared to rEndoA1 ΔH0. The number of worms analyzed for each genotype is indicated in the bar graphs. Error bars represent SEM. Scale bars indicate 2 µm.

To determine whether the function of endophilin H0 *in vivo* could be replaced by other curvature-sensing amphipathic motifs, we examined locomotion rates of transgenic *unc-57 endophilin* mutant worms expressing various rEndoA1 chimeras (Fig.2C-D). We found that several of these rEndoA1 chimeras significantly recovered the locomotion defects in *unc-57 endophilin* mutants. Among these, amphipathic motifs from BIN1 (Bridging Integrator 1, also known as Amphiphysin-2; (Low *et al*., 2008)), NUP133 (Nucleoporin 133, a component of the nuclear pore complex; (Doucet et al., 2010)), and Nadrin2 (Rho GTPase activating protein 17; (Galic et al., 2012)) significantly restored rEndoA1 H0 function. And yet other amphipathic motifs, including those from synaptic vesicle binding proteins (*e.g.*, complexin CPX-1 (Snead et al., 2014; Wragg et al., 2013), α-Synuclein (Davidson et al., 1998; Georgieva et al., 2008; Jao et al., 2008)) and intracellular membrane trafficking (*e.g.*, ARF1 (Goldberg, 1998), KES1 (Im et al., 2005; Moser von Filseck et al., 2015), and Sar1p (Lee *et al*., 2005)), either failed or had only minor activity in substituting rEndoA1 *in vivo*. These results demonstrate that endophilin H0 could carry a level of functional specificity that is only shared by a subset of curvature-sensing amphipathic motifs. Interestingly, such functional specificity could not be extrapolated from the cellular activity or localization of the parent proteins harboring the amphipathic motifs. For example, NUP133 functions at nuclear pores (Doucet *et al*., 2010; Lutzmann et al., 2002; Nordeen et al., 2020), but its amphipathic motif could replace the function of rEndoA1 H0 at synapses. Together, these data suggest that additional mechanisms, beyond sensing membrane curvature, need to be engaged for defining *in vivo* specificity of amphipathic motifs.

### Two ALPS motifs show drastic differences in their ability to replace rEndoA1 H0 at synapses

To understand the functional specificity of amphipathic motifs, we focused on two chimeric proteins: NUP133_rEndoA1 and KES1_rEndoA1. Biochemical studies have shown that the amphipathic motifs of NUP133 and KES1 act in a similar way to sense membrane curvature *in vitro*, and thereafter recognized them as two founding members of the family of ALPS motifs (Drin *et al*., 2007). Despite the biochemical similarity, NUP133 and KES1 ALPS motifs exhibited drastic differences in their ability to replace rEndoA1 H0 *in vivo* (Fig.2D). Expression of NUP133_rEndoA1 in *unc-57 endophilin* mutants significantly improved locomotion rates (with NUP133_rEndoA1 worms moving at ∼127 µm/s, vs. *unc-57* mutants at ∼18 µm/s). By contrast, worms carrying the KES1_rEndoA1 chimera had locomotion rates of ∼44 µm/s, which was significantly less than that of NUP133_rEndoA1 worms.

One possible reason for the functional deficiency of KES1_rEndoA1 was that this chimeric protein failed to localize to synapses. To examine this possibility, we compared the synaptic and axonal distribution of the proteins rEndoA1 wild type (wt), the H0-truncated mutant (ΔH0), NUP133_rEndoA1, and KES1_rEndoA1, which were all tagged with mScarlet (a red fluorescent protein) (Fig.2E). We found that fluorescently-labeled wild type endophilin was highly enriched in synaptic puncta. In contrast, the fluorescence intensity of the ΔH0 mutant puncta was significantly reduced, as previously reported (Bai *et al*., 2010) (Fig.2E). The synaptic and axonal distribution of both NUP133_rEndoA1 and KES1_rEndoA1 proteins was identical to that of wild type rEndoA1 (Fig.2E). These data show that the KES1_rEndoA1 was trafficked to synapses and enriched there, yet it could not fully replace the function of endophilin. Therefore, mechanisms other than protein-trafficking defects are needed to explain the functional difference between NUP133_rEndoA1 and KES1_rEndoA1.

To define the synaptic activity of NUP133_rEndoA1 and KES1_rEndoA1, we examined the levels of synaptic transmission using electrophysiological recordings. Due to their endocytosis defects, *unc-57 endophilin* mutants have a smaller pool of synaptic vesicles and a corresponding decrease in synaptic transmission (Bai *et al*., 2010; Kittelmann *et al*., 2013; Schuske *et al*., 2003). Specifically, *unc-57 endophilin* mutants had a decreased rate of endogenous EPSCs at neuromuscular junctions (Fig.3A, endogenous EPSC rates: wt 53 ± 3 Hz, *unc-57* 11 ± 1 Hz; p < 0.001), while the mean amplitude of endogenous EPSCs was not altered. The amplitude of evoked EPSCs in *unc-57* mutants was also significantly reduced to 0.95 ± 0.1 nA, compared to 3.2 ± 0.2 nA in wild type worms (Fig.3B; p < 0.001). These defects of synaptic transmission in *unc-57 endophilin* mutants were fully rescued upon expression of chimeric NUP133_rEndoA1 (Fig.3A, NUP133_rEndoA1 endogenous EPSC rate: 64 ± 4 Hz, and Fig.3B, evoked EPSC amplitude: 3.9 ± 0.2 nA), demonstrating a functional similarity between the NUP133 ALPS motif and rEndoA1 H0. In comparison, KES1_rEndoA1 showed only partial activity for improving synaptic transmission in *unc-57 endophilin* mutants (Fig.3A, KES1_rEndoA1 endogenous EPSC rate: 26 ± 2 Hz, and Fig.3B, evoked EPSC amplitude: 1.6 ± 0.2 nA). Together, these results demonstrate that for sustaining synaptic transmission *in vivo*, the NUP133 ALPS is a more efficient substitute for the role of endophilin H0 motif than the KES1 ALPS.

**Figure 3.**
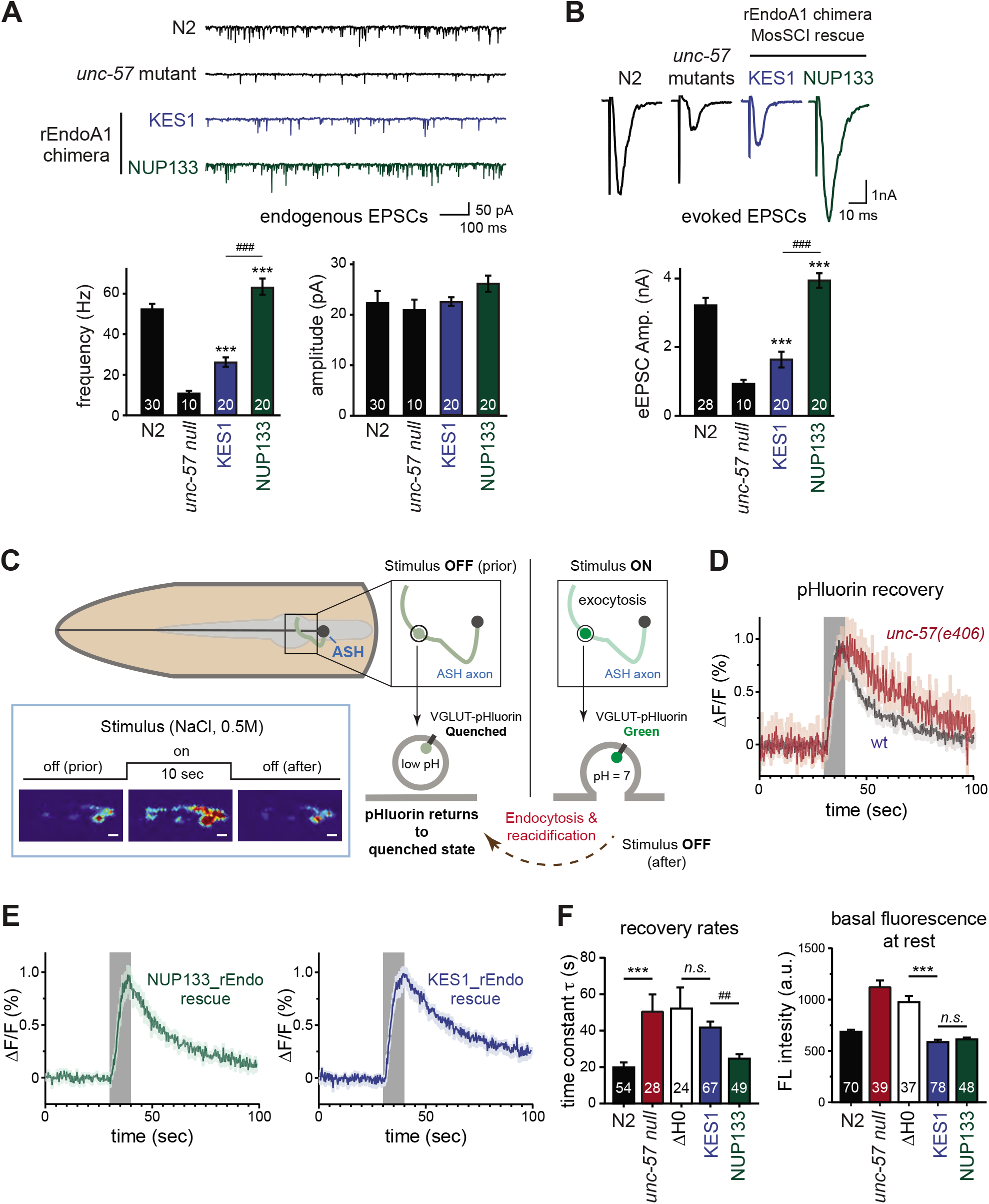
The KES1_rEndoA1 chimera exhibited reduced activity in supporting synaptic transmission and recycling synaptic vesicle proteins. Single-copy transgenes that encode GFP-tagged rEndoA1 chimera with KES1 and NUP133 amphipathic motifs (KES1_rEndo and NUP133_rEndo) were introduced into *endophilin unc-57(e406)* mutant worms. **(A-B)** Synaptic transmission was determined using electrophysiological recordings at neuromuscular junctions. Representative traces and summary data for endogenous EPSC frequency and amplitude were shown in **(A)**, and evoked EPSC (eEPSC) traces and summary data were shown in **(B)**. Schemes shown in **(C)** illustrate *in vivo* analysis of VGLUT-pHluorin fluorescence in *C. elegans* sensory neurons (ASH) upon sensory stimulation. VGLUT-pHluorin; a pH-sensitive super-ecliptic pHluorin reporter, was inserted into the first lumenal domain of the vesicular glutamate transporter VGLUT EAT-4 (Ventimiglia and Bargmann, 2017). Transgenic worms expressing VGLUT-pHluorin in the sensory ASH neuron were challenged for 10 seconds by 0.5M NaCl (“stimulus ON”). Exocytosis of synaptic vesicles triggered by NaCl stimulation delivers VGLUT-pHluorin to the plasma membrane, causing an increase in the pHluorin fluorescence in ASH neurons. After removal of NaCl stimuli (“stimulus OFF”), surface VGLUT-pHluorin was retrieved from the plasma membrane via endocytosis. Reacidification of synaptic vesicles quenches the pHluorin fluorescence. Representative images of VGLUT-pHluorin that were acquired before, during, and after NaCl stimulation were shown in the insert. Scale bars indicate 2 µm. Data plotted in **(D)** indicated mean VGLUT-pHluorin responses from wild type and *endophilin unc-57(e406)* mutant worms. Data plotted in **(E)** indicated mean VGLUT-pHluorin responses from transgenic *unc-57(e406)* mutant worms carrying KES1_rEndo *(left)* and NUP133_rEndo *(right)*. Grey rectangles indicated “stimulation-on”. Shading indicated SEM. **(F)** Quantification of the recovery time constants *(left)* and the basal levels of VGLUT-pHluorin fluorescence *(right)*. ***, p < 0.001; when compared to *endophilin unc-57 mutants*. ^###^, p < 0.001 and ^##^, p < 0.01; when compared to KES1_rEndo chimera. The number of synapses analyzed for each genotype is indicated in the bar graphs. Error bars indicate SEM.

To determine the impact of NUP133_rEndoA1 and KES1_rEndoA1 on synaptic vesicle recycling, we used imaging analysis to follow the release and retrieval of synaptic vesicles from the glutamatergic neuron ASH in intact worms (Ventimiglia and Bargmann, 2017). As shown in Fig.3C, a pH-sensitive GFP (super-ecliptic pHluorin) was inserted into the first lumenal domain of the vesicular glutamate transporter EAT-4 (VGLUT), resulting in a fluorescent VGLUT-pHluorin reporter. At the resting state, VGLUT-pHluorin fluorescence was quenched by the acidic pH of the vesicle lumen. Synaptic vesicle exocytosis delivered VGLUT-pHluorin to the plasma membrane, resulting in an increase of green fluorescence due to de-quenching of pHluorin at neutral extracellular pH. Subsequently, endocytosis and reacidification took place, returning VGLUT-pHluorin back to the quenched state. As expected, *unc-57 endophilin* mutants had significantly slowed VGLUT-pHluorin retrieval after stimulus removal (Fig.3D and 3F left panel), showing that the speed of synaptic vesicle recycling is impaired in neurons lacking endophilin. In addition, baseline VGLUT-pHluorin fluorescence in the ASH neuron was higher in *unc-57 endophilin* mutants than in wild type worms (Fig.3F right panel), indicating that an elevated level of VGLUT-pHluorin remained on the cell surface at steady-state. These results were consistent with previous reports showing the importance of endophilin for recycling synaptic vesicles (Bai *et al*., 2010; Dickman *et al*., 2005; Kittelmann *et al*., 2013; Kroll *et al*., 2019; Milosevic *et al*., 2011; Schuske *et al*., 2003; Verstreken *et al*., 2002; Watanabe *et al*., 2018). Expression of NUP133_rEndoA1 fully recovered the retrieval rates (Fig.3E left panel and Fig.3F left panel) and the basal levels of surface VGLUT-pHluorin in *unc-57* mutants (Fig.3F right panel). Conversely, expression of KES1_rEndoA1 failed to accelerate the VGLUT-pHluorin retrieval, *i.e.*, the time constant in KES1_rEndoA1 neurons was significantly higher than that in wild type or NUP133_rEndoA1 neurons (Fig.3E right panel and Fig.3F left panel), and similar to that in neurons carrying rEndoA1_ ΔH0 (Fig.3F left panel). These data demonstrate that KES1_rEndoA1 is less efficient than NUP133_rEndoA1 in supporting the retrieval of synaptic vesicle protein VGLUT-pHluorin. However, the basal level of surface VGLUT-pHluorin returned to the wild-type level upon expression of either KES1_rEndoA1 or NUP133_rEndoA1, suggesting that KES1_rEndoA1 could mediate the retrieval of VGLUT-pHluorin from the cell surface, although it is slower and less efficient.

### Synapses rescued by the KES1_rEndo chimera have abnormal endosome-like structures

To better understand the deficiency of KES1_rEndoA1, we performed high-pressure freeze electron microscopy to examine synapses at the ultrastructural level (Fig.4A). Our results identified several abnormal features at KES1_rEndoA1 synapses. First, the abundance of synaptic vesicles was only partially restored in transgenic *unc-57 endophilin* mutants that carried KES1_rEndoA1 (Fig.4A and 4B left panel). The number of synaptic vesicles for each synaptic profile was 39 ± 2 for wild type, 16 ± 1 for *unc-57* mutants, and 26 ± 1 for KES1_rEndoA1 worms (Fig.4B left panel). These ultrastructural data showing a smaller pool of the synaptic vesicles at KES1_rEndoA1 synapses were consistent with the electrophysiology findings of reduced synaptic transmission (Fig.3A-B). Second, more large clear vesicles (50-100 nm diameter) appeared at KES1_rEndoA1 synapses (Fig.4C left panel). Synaptic vesicles and large clear vesicles also became larger at KES1_rEndoA1 synapses (Fig.4B right panel for synaptic vesicles, and Fig.3C right panel for large clear vesicles) than those in wild type or *unc-57* mutant synapses. Because the amplitude of endogenous EPSCs remained unchanged at KES1_rEndoA1 synapses, the enlarged vesicles were likely deficient for releasing neurotransmitters. Third, we found a significant increase in the number of endosome-like structures (>100 nm in diameter) at KES1_rEndoA1 synapses (0 for wild type, 0.4 ± 0.1 per synapse in *unc-57* mutants, and 1.5 ± 0.2 per KES1_rEndoA1 synapse; Fig.4D left panel). Moreover, as shown in the Fig.4D right panel, the endosome-like structures occurred more frequently at KES1_rEndoA1 synapses (16 out of *(i)* 18) than at *unc-57* mutant synapses (9 out 25). These increases in abundance and frequency of endosome-like structures at KES1_rEndoA1 synapses were surprising, and raised questions regarding the source of membranes for producing the endosome-like structures. Because our results showed that KES1_rEndoA1 synapses had fewer synaptic vesicles but more endosome-like structures, we examined the possibility that endosome-like structures and synaptic vesicles compete for membranes. Consistent with this idea, we found an inverse correlation between the synaptic vesicle abundance and the size of endosome-like structures at synapse (Fig.4E), supporting a competitive nature between these two distinct types of membrane organelles.

**Figure 4.**
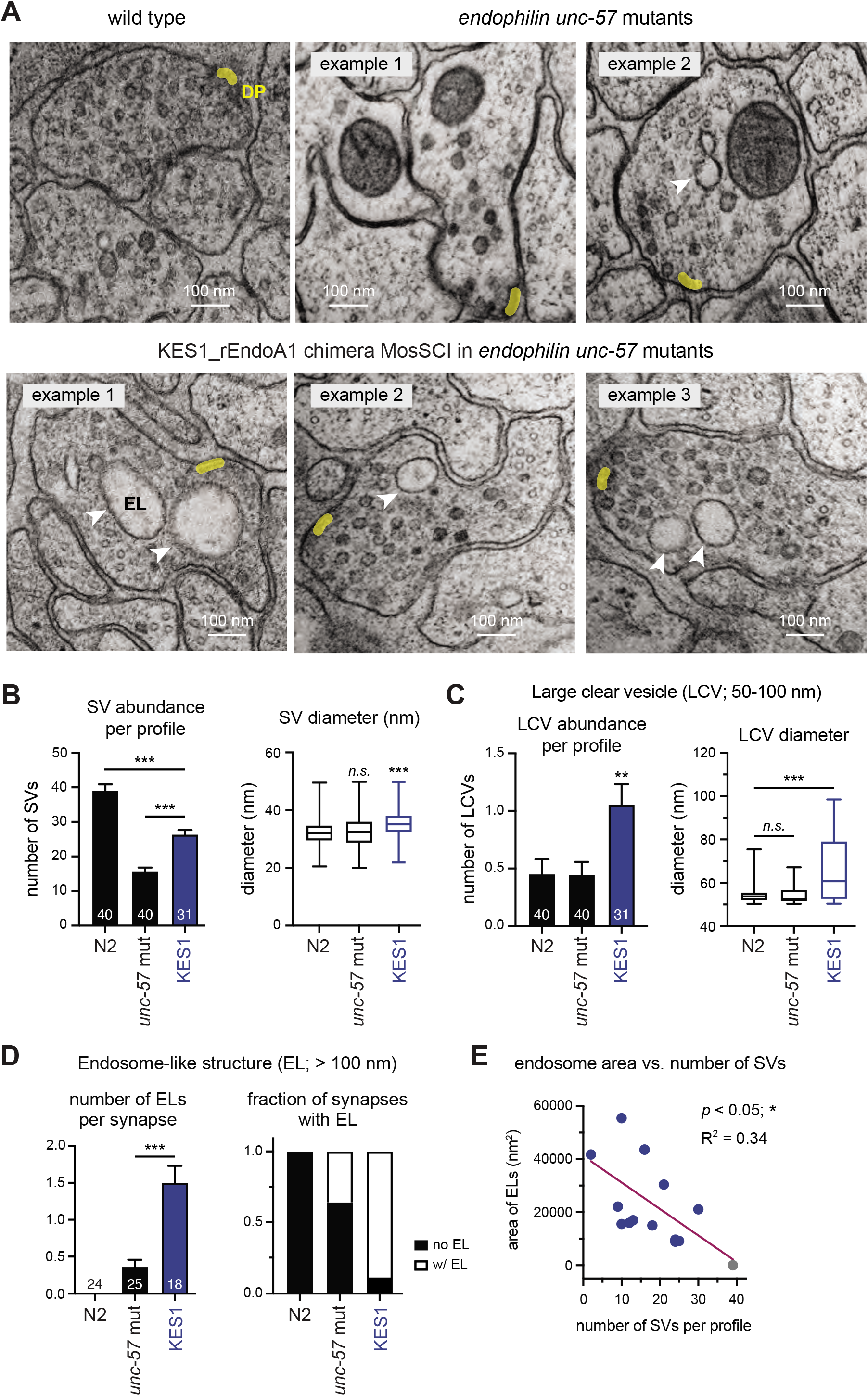
Synapses rescued by the KES1_rEndo chimera have increased accumulation of abnormal endosome-like structures. Single-copy transgenes that encode GFP-tagged KES1_rEndoA1 chimera were introduced into *endophilin unc-57(e406)* mutant worms. **(A)** Representative transmission electron micrographs of synapses in the ventral nerve cords of adult hermaphrodites. Yellow shading indicates dense projections (DP). White arrow heads indicate endosome like structures (EL; >100 nm in diameter). Scale bars indicate 100 nm. **(B)** Quantification of synaptic vesicles (SV; <50 nm in diameter). The abundance of SVs per synaptic profile was shown as mean ± standard error in the *left* panel. Error bars indicate SEM. The diameter of SVs was presented as box-whisker plots in the *right* panel. Whiskers indicate minimum and maximum values. **(C)** Quantification of large clear vesicles (LCV; 50-100 nm in diameter). The abundance of LCVs per synaptic profile was plotted as mean ± standard error in the *left* panel. Error bars indicate SEM. The diameter of LCVs was presented as box-whisker plots in the *right* panel. Whiskers indicate minimum and maximum values. Data for SV and LCV abundance and size were collected from 40 wild type, 40 *unc-57* mutant, and 31 KES1_rEndoA1 synaptic profiles. **(D)** Endosome-like structures (EL; >100 nm in diameter). The number of ELs per synapse was presented as mean ± standard error (*left*). Error bars indicate SEM. ***, p < 0.001; **, p < 0.01; n.s., not significant; one-way ANOVA followed by Dunnett’s test. The fraction of synapses with and without endosome-like structures was calculated and plotted for each genotype (*right*). Data for endosome-like structures were collected from 24 wild type synapses, 25 *unc-57* mutant synapses, and 18 KES1_rEndoA1 synapses. **(E)** A scatter plot illustrated the negative relationship between the size of endosome-like structures and the abundance of synaptic vesicles. The y-axis value of each dot indicates the sum of maximum cross-sectional areas of endosome-like structures in a synapse. The corresponding x-axis value indicates the mean number of SVs per synaptic profile of the same synapse where the y-value was determined. Data obtained from KES1_rEndo synapses were colored in blue. The grey dot indicates data from wild type synapses (the mean number of synaptic vesicles per profile and the lack of endosome-like structures). The purple line indicated the best fit of simple linear regression. R^2^ = 0.34 and * p < 0.05. Statistical analysis and graphing were carried out using Prism 9 (GraphPad).

Next, we sought to determine factors that limit the ability of the KES1 ALPS motif to replace endophilin H0. We reasoned that if we could identify strategies to improve the function of KES1_rEndoA1 at synapses, the resulting knowledge would lead to a deeper understanding of how endophilin H0 works *in vivo*. To this end, we first looked into biochemical differences among endophilin H0, NUP133 ALPS, and KES1 ALPS motifs. It was previously suggested that the abundance of basic residues is a major factor that differentiates endophilin H0 from members in the ALPS family (Drin and Antonny, 2010; Drin *et al*., 2007). Interestingly, while the ALPS motifs of NUP133 and KES1 both belong to the ALPS family (Drin *et al*., 2007), they have different cationic levels. As shown in Fig.5A, the KES1 ALPS motif has a net charge level of 0, and the NUP133 ALPS motif has a net charge level of +2, whereas rEndoA1 H0 has a net charge level of +4. Alignment of amphipathic helix sequences showed that the net charge levels of KES1, NUP133, and rEndoA1 motifs remained largely constant across orthologs from various species (Fig.S3), suggesting that the levels of positive charges of these motifs are preserved during evolution.

**Figure 5.**
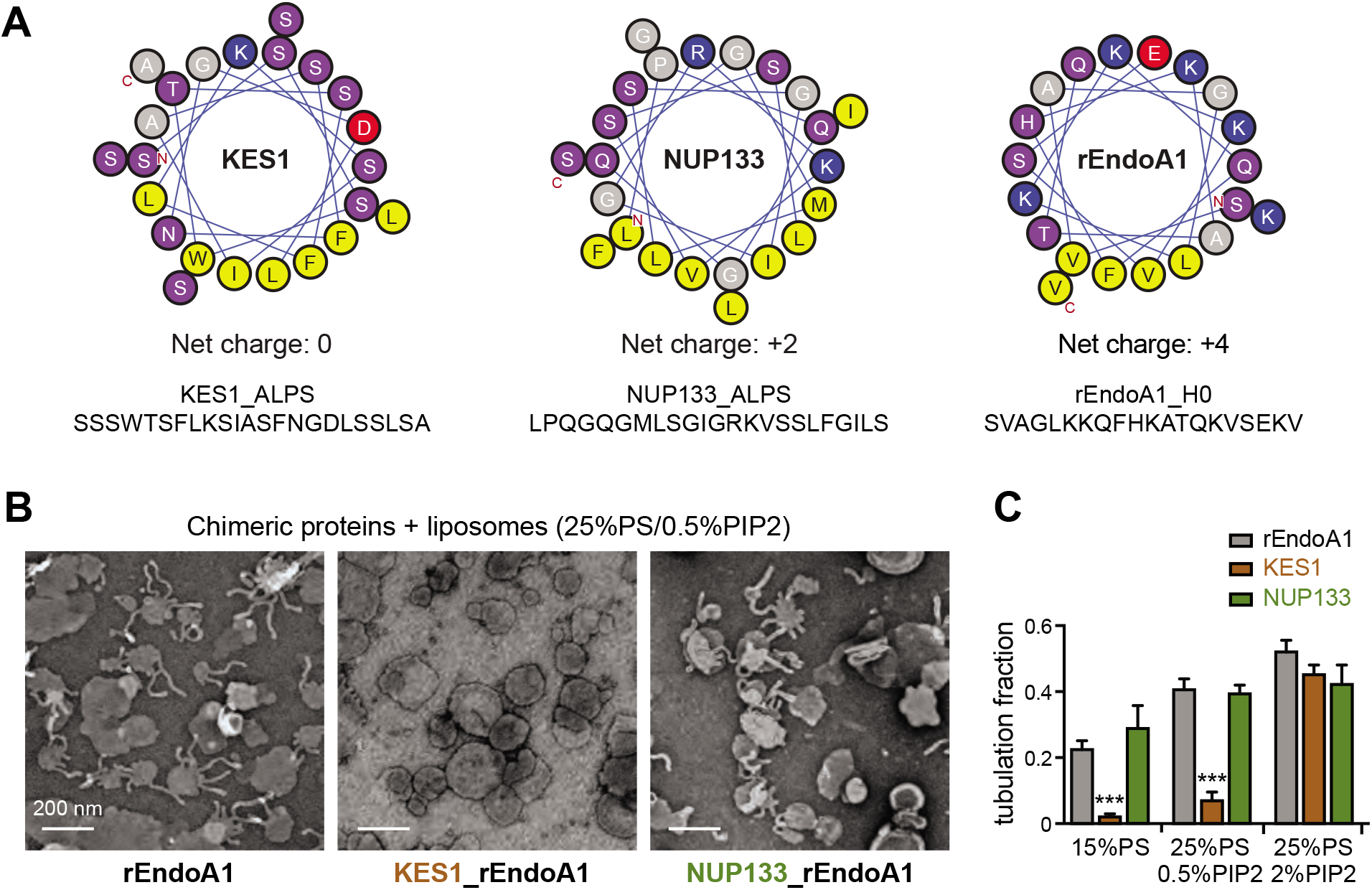
Chimeric endophilin proteins carrying KES1 and NUP133 amphipathic helixes display distinct properties. **(A)** Helical wheel presentations (top) and sequences (bottom) of amphipathic helices in KES1, NUP133, and rEndoA1. ALPS: <u>a</u>mphipathic lipid <u>p</u>acking <u>s</u>ensor motif. Net electric charges of these amphipathic helices were shown (middle). Amino acid residues in the helical wheel presentations are colored as described in the legend of Figure 1. **(B)** Transmission electron micrographs for liposomes incubated with wild type and chimeric rEndoA1 proteins; *(left)* wild type rEndoA1, *(middle)* KES1_rEndoA1, and *(right)* NUP133_rEndoA1. Liposomes were prepared by extrusion through filters with 0.2 µm pore size. The lipid composition of these liposomes was 25%PS/0.5%PIP2/74.5%PC. Wild type and chimeric rEndoA1 proteins (4µM) were incubated with liposomes (2 mM total lipids) for 5 min at the room temperature. Reaction was stopped by fixing with 0.5x Karnovsky’s solution. Scale bars: 200 nm. **(C)** Quantification of membrane tubulation. Tubulation Fraction = (# of liposomes with extended protrusions) / (# of total liposomes). Liposomes contained either 15%PS/85%PC, 25%PS/0.5%PIP2/74.5%PC, or 25%PS/2%PIP2/73%PC, respectively. Error bars indicate SEM. ***, p < 0.001, compared to liposomes incubated with wild type rEndoA1. One-way ANOVA followed by Dunnett’s test was used for statistical analysis.

### The charge level of KES1_ALPS limits its potential to substitute endophilin H0 in vitro

To determine how grafted amphipathic motifs influence chimeric rEndoA1, we used electron microscopy to study the ability of chimeric rEndoA1 to generate membrane tubules. Previous studies showed that the membrane tubulation activity of wild type endophilin requires interactions with lipids of negatively charged headgroups (*e.g.*, phosphatidylserine PS and phosphatidylinositol 4,5-bisphosphate PIP2) (Farsad *et al*., 2001; Gallop *et al*., 2006; Jao et al., 2004; Masuda *et al*., 2006). Consistent with these early findings, our control experiments showed that wild type rEndoA1 generated more, and longer, membrane tubules on liposomes with correspondingly increased amounts of negatively charged lipids (Fig.S4). When we compared NUP133_rEndoA1 and wild type rEndoA1, we found that these two proteins generated identical amounts of membrane tubules on liposomes (Fig.5B and 5C). By contrast, the KES1_rEndoA1 chimera failed to produce abundant membrane tubules on liposomes carrying either 15%PS/85%PC or 25%PS/0.5%PIP2/74.5%PC (Fig.5B and 5C). The reduction in membrane tubulation by KES1_rEndoA1 was likely due to changes of electrostatic sensing, rather than the membrane tubulation capability *per se*, because KES1_rEndoA1 was able to robustly produce membrane tubules when membrane PIP2 was increased to 2% (Fig.5C). Together, these data show that KES1_rEndoA1 requires a higher amount of negatively charged lipids to tubulate liposomes (PIP2 bears four negative charges whereas PS has one), raising a possibility that the net charge level of 0 of KES1 ALPS limits the potential of this motif to substitute endophilin H0.

### Increasing net positive charges of the KES1 ALPS motif improves its synaptic function

To define the relationship between the charge distribution and the functionality of amphipathic motifs, we engineered mutations to improve the function of the KES1_rEndoA1 chimera *in vivo*. If the net charge level was indeed a limiting factor, we reasoned that introducing positive charges into the KES1 ALPS should improve its functionality. As a member of the ALPS family, the amphipathic motif of KES1 is rich in hydrophilic residues such as serine (S) and threonine (T). To add positive charges to the KES1 ALPS motif, we introduced mutations to convert hydrophilic S/T residues into positively charged lysine (K) residues. As shown in Fig.6A, we generated single K mutations to S2, S3, and S4, which are three residues in the first turn of the KES1 amphipathic helix. We found that these single mutations (S2K, S3K, or S4K) significantly improved the activity of KES1_rEndoA1 to restore locomotion rates in *unc-57 endophilin* mutants (Fig.6B and 6C). Interestingly, S2K, S3K, or S4K mutations led to a similar level of locomotory improvement, suggesting that the position of these mutations had little or no impact on the functional changes of mutant KES1_rEndoA1. Indeed, when we systematically introduced single K mutations to change each S/T site in the KES1 motif, we found that every single S/K mutant protein partially, but significantly, restored locomotion activity, and the locomotion rates were similar among the KES1_rEndoA1 S/K mutant worms, regardless of the position of the mutation (Fig.6C). These data suggest that the S/K mutations improved KES1_rEndoA1 function by increasing the net positive charge rather than by altering specific charge locations.

**Figure 6.**
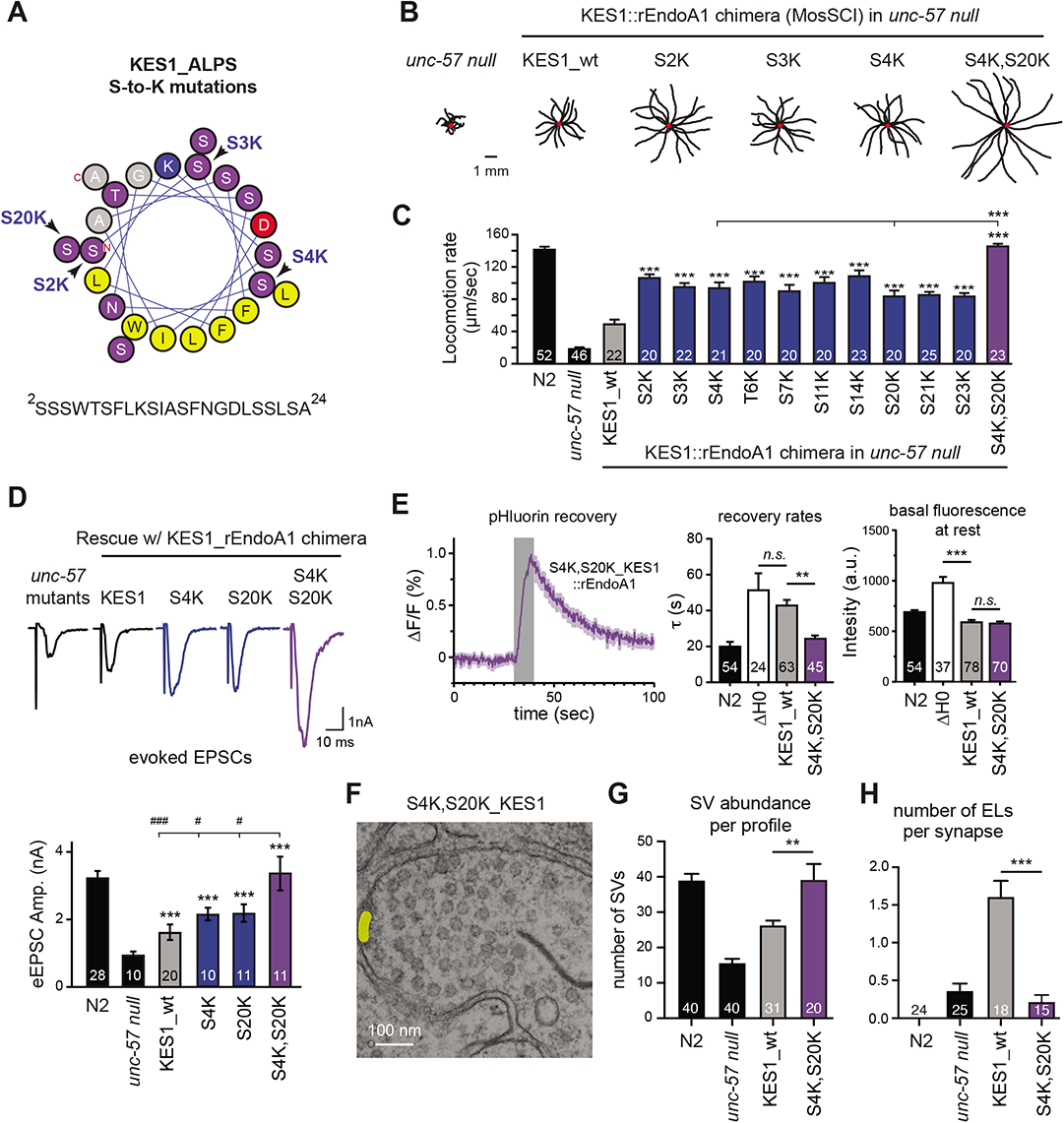
Serine to Lysine mutations improved the function of KES1_rEndoA1 chimera *in vivo*. A helical wheel presentation of the ALPS motif of KES1 was shown in **(A)**. Arrows indicated the serine residues (S2, S3, and S4) that occupy the 1^st^ turn of the amphipathic helix, and S20 that resides on the opposite side to S4. The full sequence of the ALPS motif of KES1 was listed below the helical wheel. **(B)** Representative trajectories (15 animals) of 30s locomotion were shown for transgenic *endophilin unc-57* mutant worms carrying various KES1_rEndoA1 chimeras as indicated. Locomotion traces of *endophilin unc-57* mutant worms were shown as controls. The scale bar indicates 1 mm. **(C)** Quantification of locomotion rates. Mutant KES1_rEndoA1 chimeras with serine-to-lysine (S/K) mutations were examined for their ability to restore locomotion activity in *endophilin unc-57* mutant worms. The number of worms analyzed for each genotype is shown in the bar graphs. Error bars represent SEM. **(D)** Representative eEPSC (evoked EPSC) traces and summary data for eEPSC amplitudes were shown for indicated genotypes. ***, p < 0.001; when compared to *endophilin unc-57 mutants*. ^###^, p < 0.001; and ^#^, p < 0.05; when compared to animals rescued by S4K,S20K mutant KES1_rEndoA1. The number of worms analyzed for each genotype is indicated in the bar graphs. Error bars indicate SEM. One-way ANOVA followed by Dunnett’s test was used for statistical analysis. Data plotted in **(E)** showed VGLUT-pHluorin responses from worms of indicated genotypes - *(left) a* time course of mean fluorescence responses in ASH neurons carrying S4K,S20K KES1_rEndoA1, *(middle)* rate constants of fluorescence decay; and *(right)* basal levels of VGLUT-pHluorin fluorescence. Grey rectangles indicated “stimulation-on”. Shading indicated SEM. Error bars in the middle and right panels indicated SEM. The number of synapses analyzed for each genotype was indicated in the bar graphs. **(F)** A representative transmission electron micrograph of synapses rescued by the S4K,S20K KES1_rEndoA1. **(G)** The abundance of SVs per synaptic profile was shown as mean ± standard error. Data for SV abundance were collected from 40 wild type, 40 *unc-57* mutant, 31 KES1_rEndoA1 and 20 S4K,S20K KES1_rEndoA1 synaptic profiles. Error bars indicated SEM. **(H)** The number of endosome-like (EL) structures per synapse was presented as mean ± SEM. Data for endosome-like structures were collected from 24 wild type synapses, 25 *unc-57* mutant synapses, 18 KES1_rEndoA1 synapses, and 15 S4K,S20K_KES1_rEndoA1 synapses. ***, p < 0.001; **, p < 0.01; n.s., not significant; one-way ANOVA followed by Dunnett’s test.

Since single S/K mutations only partially restored the locomotion activity, we asked whether addition of a second K residue, bringing the net charge from +1 to +2, would further improve the activity of KES1_rEndoA1 *in vivo*. Indeed, a combination of two single S/K mutations we tested (S4K and S20K, Fig.6C) showed significantly enhanced activity in restoring locomotion of *unc-57 endophilin* mutant worms, when compared to the single S4K and S20K mutants (Fig.6B and 6C). Next, we determined the impact of S4K,S20K KES1_rEndoA1 at synapses using both functional and ultrastructural analyses. First, electrophysiological recordings showed that expression of S4K,S20K KES1_rEndoA1 fully restored the amplitude of evoked EPSCs (Fig.6D), and the frequency of endogenous EPSCs (Fig.S5) in *unc-57 endophilin* mutants. Second, we found that VGLUT-pHluorin retrieval at synapses was fully recovered by expression of S4K,S20K KES1_rEndoA1 in *unc-57 endophilin* mutants (Fig.6E). Third, electron microscopy results showed that the abundance of synaptic vesicles at S4K,S20K KES1_rEndoA1 synapses was recovered to a level close to that in wild type synapses (Fig.6F and 6G). Further, the number of endosome-like structures was significantly reduced, when compared to synapses carrying KES1_rEndoA1 (Fig.6H). Together, these findings demonstrate that increasing the net charge to +2 improved the activity of KES1_rEndoA1 enough to substitute endophilin H0, and enabled mutant KES1_rEndoA1 to efficiently support synaptic activities.

### The synaptic function of DivIVA_rEndoA1 is improved by introducing positive net charges into its curvature-sensing motif

To determine how other amphipathic motifs respond to changes of net charge level, we investigated the sequence from DivIVA, a bacterial protein involved in cell division (Hempel et al., 2008; Lenarcic et al., 2009). The DivIVA amphipathic motif shares a couple of similar properties with the KES1 ALPS motif. First, expression of the DivIVA_rEndoA1 chimeric protein failed to restore locomotion activity in *unc-57 endophilin* mutant worms (Fig.2C and 2D). Second, the net charge level of the DivIVA amphipathic motif is 0 (Fig.7A), which is identical to that of KES1 ALPS. However, unlike KES1 ALPS, which lacks a significant amount of charged residues, the DivIVA amphipathic motif has 3 positive charges (from Lys and Arg) and 3 negative charges (from Asp and Glu), resulting in the net charge of 0 (Fig.7A). This unique electrostatic property allowed us to alter the net charge level in another way. Instead of using K residues, we introduced E4S, D12S, and E14S mutations into the DivIVA amphipathic motif to bring the net charge level to +3. We then systematically examined the *in vivo* function of the wild type (wt) and triple mutant (E4S,D12S,E14S) versions of DivIVA_rEndoA1 chimeras using behavioral, electrophysiological, imaging, and ultrastructural analyses. In contrast to the failure of restoring locomotion by wt_DivIVA_rEndoA1, expression of the triple mutant E4S,D12S,E14S_DivIVA_rEndoA1 chimera in *unc-57* mutant worms fully restored locomotion (Fig.7B), endogenous EPSCs (Fig.7C), and evoked EPSCs (Fig.7D), as well as retrieval rates and basal levels of VGLUT-pHluorin (Fig.7E). Ultrastructural data showed that expression of the wt_DivIVA_rEndoA1 chimera in *unc-57* mutant animals failed to recover the synaptic vesicle abundance and led to the formation of abnormal endosome-like structures at synapses (Fig.7F, and 7G). In contrast with those results, at synapses carrying the E4S,D12S,E14S_DivIVA_ rEndoA1 chimera, we found that the number of synaptic vesicles was returned to a normal level, and no endosome-like structures were observed (Fig.7F, and 7G). Together, these data present a second case where the function of endophilin chimeras could be improved *in vivo* by introducing positive net charges into a curvature-sensing motif.

**Figure 7.**
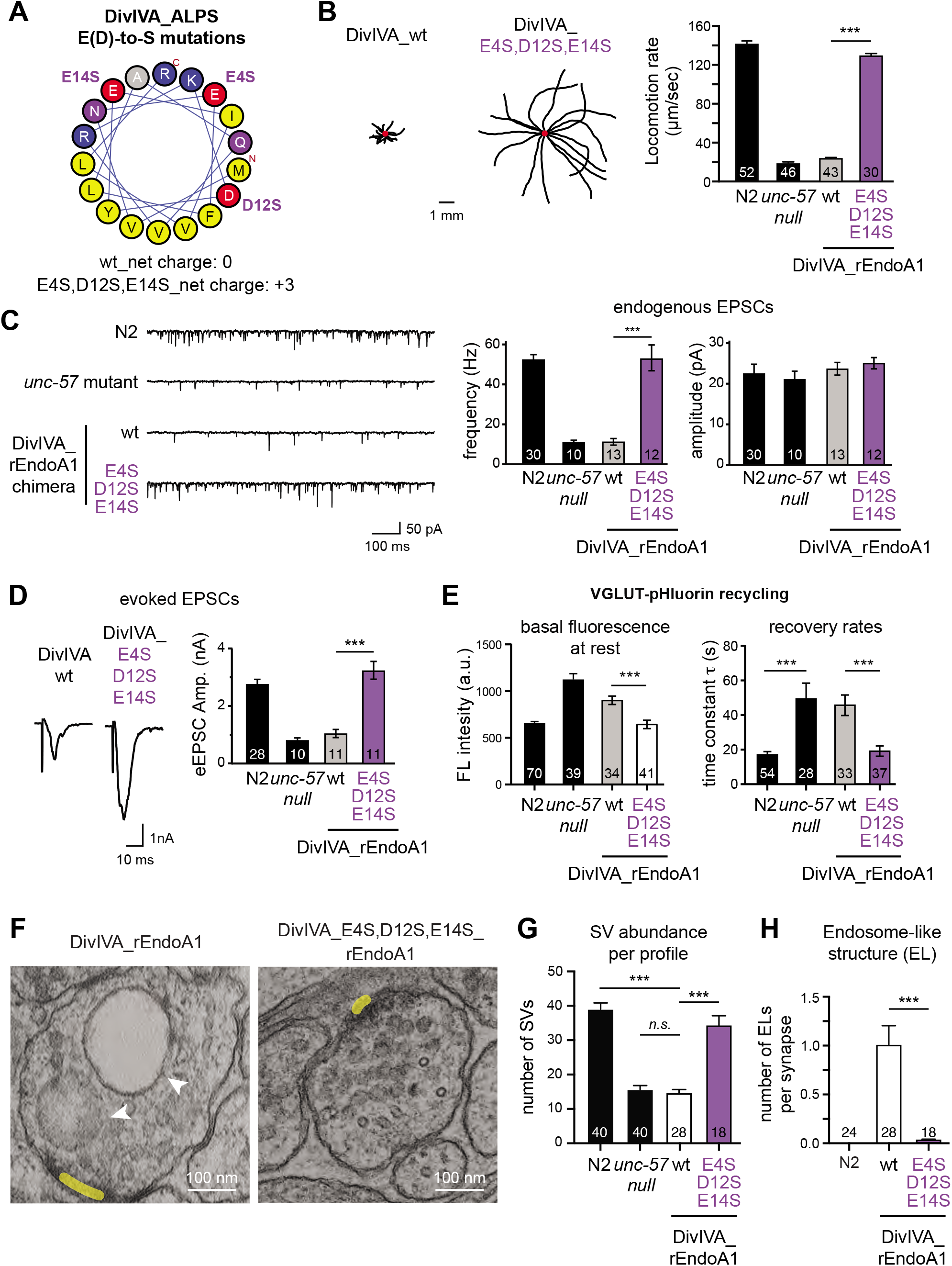
Increases of positive net charges of the amphipathic helix improved the function of DivIVA_rEndoA1 chimeras. A helical wheel presentation of an ALPS motif of the bacteria protein DivIVA was shown in **(A)**. Two glutamic acid residues (E4, E14) and an aspartic acid residue (D12) were changed to serine residues to generate the E4S,D12S,E14S version of DivIVA_rEndoA1. Data from locomotion studies **(B)**, electrophysiological recordings of endogenous EPSCs **(C)** and evoked EPSCs **(D)**, and fluorescence analysis of ASH VGLUT-pHluorin fluorescence responses to sensory stimuli **(E)**, were quantified. These results were plotted as described in the legends of Figures 3 and 6. Error bars represented SEM. The number of synapses analyzed for each genotype is indicated in the bar graphs. **(F)** Representative transmission electron micrographs of synapses rescued by DivIVA_rEndoA1 and the mutant E4S,D12S,E14S DivIVA_rEndoA1 were shown as indicated. Yellow shading indicated dense projections (DP). The white arrowheads show endosome-like structures. **(G)** The abundance of synaptic vesicles per synaptic profile was shown as mean ± SEM. Data for synaptic vesicle abundance were collected from 40 wild type, 40 *unc-57* mutant, 28 DivIVA_rEndoA1 and 18 E4S,D12S,E14S_DivIVA_rEndoA1 synaptic profiles. **(H)** The number of endosome-like (EL) structures per synapse was presented as mean ± standard error. Data for endosome-like structures were collected from 24 wild type, 25 *unc-57* mutant, 28 DivIVA_rEndoA1 and 18 E4S,D12S,E14S_DivIVA_rEndoA1 synapses. ***, p < 0.001; one-way ANOVA followed by Dunnett’s test.

### The net positive charge of the H0 motif is essential for the function of endophilin at synapses

Finally, we examined the electrostatic property of endophilin H0 and its role in supporting synaptic activity. The H0 motif of rEndoA1 has a net charge level of +4, with 5 positively charged Lys residues and 1 negatively charged Glu residue (Fig.8A). Alignment analysis showed that all Lys residues are highly conserved among orthologs of endophilin A1 across species (Fig.S3). To determine if conserved Lys residues are required, we introduced single K/S mutations to the rEndoA1 H0 motif and expressed mutant versions of rEndoA1 in *unc-57 endophilin null* worms. Interestingly, single K/S mutations had little impact on the rescue activity of rEndoA1 in restoring locomotion rates (Fig.S6A), suggesting that individual lysine residues are not essential despite being highly conserved. However, these mutations showed a clear additive effect of disrupting the function of endophilin, as locomotion rates became lower when combinations of more K/S mutations were added (Fig.8B). To understand how K/S mutations alter the function of endophilin at synapses, we performed electrophysiological recordings to determine the levels of synaptic transmission. Like the locomotion results, we found that increases in the number of K/S mutations led to slower rates of endogenous EPSCs (Fig.8C), and lower amplitudes of evoked EPSCs (Fig.8D), without affecting the amplitudes of endogenous EPSCs (Fig.S6B). In particular, the introduction of 4 K/S mutations (K7S,K8S,K18S,K20S) completely abolished the ability of rEndoA1 to restore synaptic currents, indicating that positively charged Lys residues function as a collective group. Interestingly, our ultrastructural analysis showed that synapses carrying the 4 K/S mutant rEndoA1 had more endosome-like structures and fewer synaptic vesicles (Fig.8E-G), resembling features of KES1_rEndoA1 synapses. Together, these results show that the net positive charge of the H0 motif is essential for the function of endophilin at synapses, further supporting the conclusion that the level of net charge contributes to the functional specificity of curvature-sensing motifs *in vivo*.

**Figure 8.**
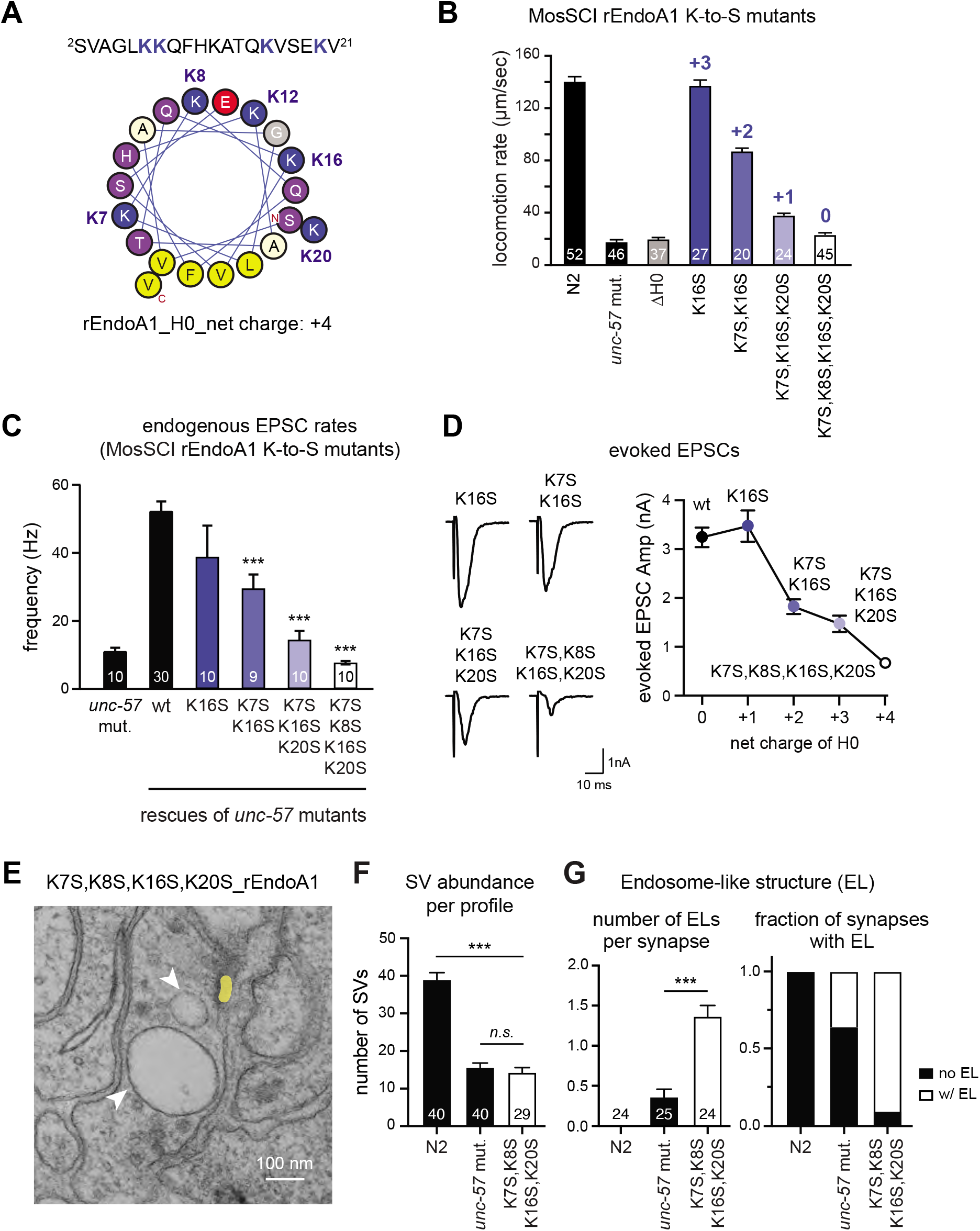
Lysine-to-serine mutations disrupted the function of Endophilin and induced accumulation of endosome-like structures at synapses. **(A)** Lysine residues were highlighted in the sequence (top) and the helical wheel presentation (middle) of the H0 motif of rEndoA1. The net charge (+4) of rEndoA1 H0 was indicated (bottom) for clarity. Single-copy transgenes that encode GFP-tagged rEndoA1 were introduced into *endophilin unc-57(e406)* mutant worms. Four versions of lysine-to-serine (K-to-S) mutant rEndoA1 (including the single K16S, the double K7S,K16S, the triple K7S,K16S,K20S, and the quadruple K7S,K8S,K16S,K20S mutants) were examined for their ability to restore locomotion **(B)**, endogenous EPSCs **(C)**, and evoked EPSCs **(D)**. Error bars represent SEM. The number of worms analyzed for each genotype is indicated in the bar graphs. **(E)** Representative transmission electron micrograph of synapses rescued by the K7S,K8S,K16S,K20S quadruple mutant rEndoA1. Yellow shading indicated dense projections (DP). White arrowheads showed endosome-like structures. **(F)** The abundance of synaptic vesicles per synaptic profile was shown as mean ± standard error. Data for SVs were collected from 40 wild type, 40 *unc-57* mutant, and 29 K7S,K8S,K16S,K20S mutant synaptic profiles. **, p < 0.01; *n.s.*, not significant; compared to the quadruple K7S,K8S,K16S,K20S mutants, one-way ANOVA followed by Dunnett’s test. **(G)** Quantification of endosome-like (EL) structures were shown. The number of ELs per synapse was presented as mean ± standard error (*left*). Error bars indicate SEM. ***, p < 0.001; one-way ANOVA followed by Dunnett’s test. The fraction of synapses with and without endosome-like structures was plotted for each genotype (*right*). Data for endosome-like structures were collected from 24 wild type synapses, 25 *unc-57* mutant synapses, and 24 K7S,K8S,K16S,K20S mutant rEndoA1 synapses.

## Discussion

Our studies examined the functional specificity of amphipathic motifs *in vivo*. First, we demonstrated that endophilin H0 could be substituted by only a subset of curvature-sensing amphipathic motifs, highlighting the functional diversity among these amphipathic motifs in neurons. Second, we found that expression of the endophilin chimera carrying the KES1 ALPS motif failed to restore synaptic function and morphology, leading to an abnormal increase of endosome-like structures and a reduced pool of synaptic vesicles at synapses. Third, we improved the synaptic function of the KES1_rEndoA1 chimera by increasing the net positive charges in the KES1 ALPS motif, demonstrating a sufficient role of electrostatic alterations in changing the functional specificity of amphipathic motifs. Fourth, we showed that adjustment of cationic status of another curvature-sensitive motif from DivIVA led to similar improvement of the synaptic activity of the DivIVA_rEndoA1 chimera, attesting to the importance of the electrostatic contribution to these two curvature-sensitive motifs from the unrelated proteins KES1 and DivIVA. Finally, we demonstrated the necessity of positive charges on the endophilin H0 motif *in vivo*. Thus, cationic property is both necessary, and sufficient, for providing curvature-sensing motifs with the functional specificity to replace endophilin H0 *in vivo*.

Amphipathic α-helical motifs are often found in proteins that regulate the trafficking of membrane organelles. These motifs are thought to carry out specific activities for assisting their parental proteins in performing distinct cellular functions. In recent years, *in vitro* studies using synthetic membranes and molecular dynamics simulations have found a wealth of plausible mechanisms for interactions between amphipathic motifs and membranes. However, due to a lack of systematic analysis *in vivo*, understanding of these motifs largely remains as working models. This study used a protein engineering approach to identify factors that are sufficient to define the specificity of amphipathic motifs *in vivo*. Our findings led to several conclusions on mechanisms governing the specificity of endophilin H0 in living neurons.

*(i)* The function of the H0 motif does not require a specific protein sequence. Our studies found that unrelated amphipathic sequences are able to substitute endophilin H0 for synaptic function, therefore uncoupling the essential function of the H0 motif from specific protein-protein interactions (*e.g.*, binding between endophilin H0 and SH3 domains (Chen *et al*., 2014; Vazquez *et al*., 2013; Wang *et al*., 2008)). In addition, the linker between the H0 motif and the BAR domain of endophilin is flexible, and it tolerates insertion of amino acid residues, indicating that H0 has structural freedom from the rest of the protein.

*(ii)* The specificity of an amphipathic motif cannot be simply extrapolated from the function of its parental protein, and instead requires investigations that include functional studies. For instance, when grafted into endophilin, the amphipathic motif from the nuclear pore complex protein NUP133 (Doucet *et al*., 2010; Lutzmann *et al*., 2002; Nordeen *et al*., 2020) can replace the endophilin H0 motif in its role at synapses. Thus, amphipathic motifs could carry similar functions, even though their parental proteins may act on different membrane organelles.

*(iii)* Electrostatic contribution is necessary and sufficient for amphipathic motifs to support the activity of endophilin at synapses. Such an electrostatic contribution appears to be independent to the location of charges, or to particular types of amino acid residues. The overall status of net charges, rather than particular amino acid residues, modulates the activity of endophilin H0. Indeed, single Lys-to-Ser mutations in the endophilin H0 motif have little impact, but combinations of the Lys-to-Ser mutations disrupt endophilin’s activity at synapses in an additive manner.

*(iv)* Endophilin H0 likely senses both curvature and electrostatic status of membranes. The amphipathic motifs of NUP133 and endophilin were previously categorized into two subgroups based on the type of residues in the polar face – the NUP133 amphipathic motif is a *bona fide* curvature sensor in the ALPS motif family (Drin and Antonny, 2010; Drin *et al*., 2007) due to abundant polar serine and threonine residues, whereas endophilin H0 is thought to have low sensitivity to membrane curvature because it is rich in charged residues. In contrast to the anticipated distinction between two subgroups of amphipathic motifs, we find that NUP133 ALPS successfully replaced endophilin H0 *in vivo*. These results suggest that the functional boundary between ALPS motifs and endophilin H0 is not absolute. It is likely that NUP133_ALPS and endophilin H0 play similar roles as dual sensors for membrane curvature and negatively charged lipids.

What is the contribution of positively charged residues in endophilin H0 to synaptic activity *in vivo*? Our results show that the cationic property of endophilin H0 has a strong impact on endosome-like structures, *i.e.*, the intermediate membrane organelles that support the endocytic reformation of synaptic vesicles (Clayton and Cousin, 2009; Hoopmann et al., 2010; Kononenko and Haucke, 2015; Watanabe et al., 2013a; Watanabe et al., 2013b; Watanabe et al., 2014). Recent studies show that, following the exocytosis of synaptic vesicles, endophilin acts at two distinct sites to recycle synaptic vesicle proteins and membranes. It promotes endocytic retrieval from the plasma membrane into endosomes (Kononenko et al., 2014; Watanabe *et al*., 2018), and then helps to resolve endosomes into synaptic vesicles (Kittelmann *et al*., 2013; Watanabe *et al*., 2018). This two-step model suggests that complete removal of endophilin from synapses would block the initial endocytic retrieval, imposing a negative impact on the formation of endosomes. Consistent with this, our results show that the probability of finding endosome-like structures at *endophilin* mutant synapses is low (Fig.4), as high levels of VGLUT-pHluorin fluorescence indicated a failure of protein retrieval from the plasma membrane at *endophilin* mutant synapses (Fig.3F). By contrast, at synapses expressing endophilin chimeras with few positive charges (*e.g.*, KES1_rEndoA1, DivIVA_rEndoA1 and rEndoA1 with 4 K/S mutations), endosome-like structures become abundant and are much more frequently observed (Fig.4, Fig.7-E-H, and Fig.8E-G). Such endosomal changes are likely at the expense of synaptic vesicles, as there is a negative correlation between the size of endosome-like structures and the abundance of synaptic vesicles (Fig.4E). Collectively, our findings indicate that endophilin requires a strong net positive charge in the H0 motif to regulate endosomes and synaptic vesicles in neurons. Amphipathic motifs with sole activities in curvature-sensing (*e.g.*, KES1_ALPS) are not sufficient for endophilin to reform synaptic vesicles from endosomes, which could lead to a ‘traffic jam’ causing abnormal accumulation of endosomes at synapses.

Results in this study provide *in vivo* evidence supporting a dual sensing capability of endophilin H0 to detect both curvature and electrostatic states of membranes. The cationic property of H0 is likely more important for the role of endophilin on endosomes than on the plasma membrane, because these membranes contain anionic lipids with distinct properties (Balla, 2013; Di Paolo and De Camilli, 2006). The plasma membrane is abundant with phosphatidylinositol 4,5-bisphosphates which bear a negative charge of ∼4, whereas endosomes are enriched for phosphatidylinositol phosphates which have a negative charge of ∼1.5. As a result, to recognize endosomal membranes, the endophilin H0 motif may require a high level of positive charges to reach sufficient electrostatic interactions. Future studies are needed to address this scenario in detail. Nonetheless, our studies demonstrate an electrostatic contribution for endophilin H0 to function *in vivo*. These results suggest a model in which the H0 motif carries a dual sensing activity of membrane curvature and electrostatic status to ensure endophilin’s action at distinct sites during synaptic vesicle endocytosis.

## Acknowledgments

This research was supported by NIH R01GM127857 and R01NS115974 to JB. The authors thank the *Caenorhabditis elegans* Genetics Center (funded by NIH Office of Research Infrastructure Programs P40 OD010440) and Dr. Cori Bargmann for worm strains. We thank Dr. Susan Parkhurst for critical reading of the manuscript. We thank the Cellular Imaging core at Fred Hutch for technical help with electron microscopy.

---

**Suppl. figure 1.**
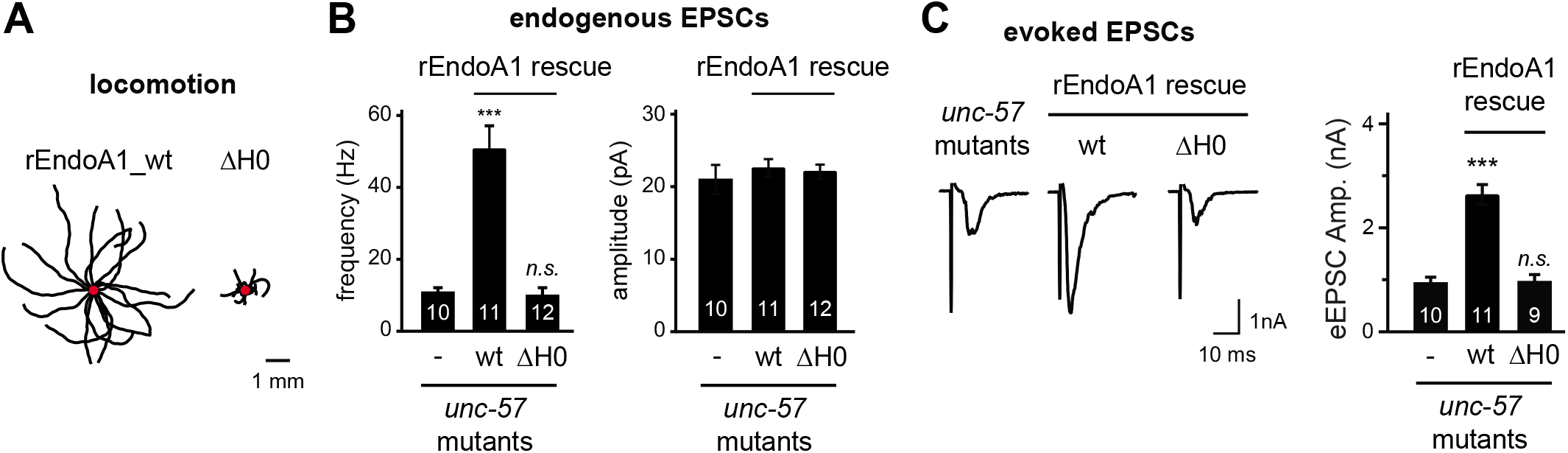

**Suppl. figure 2.**
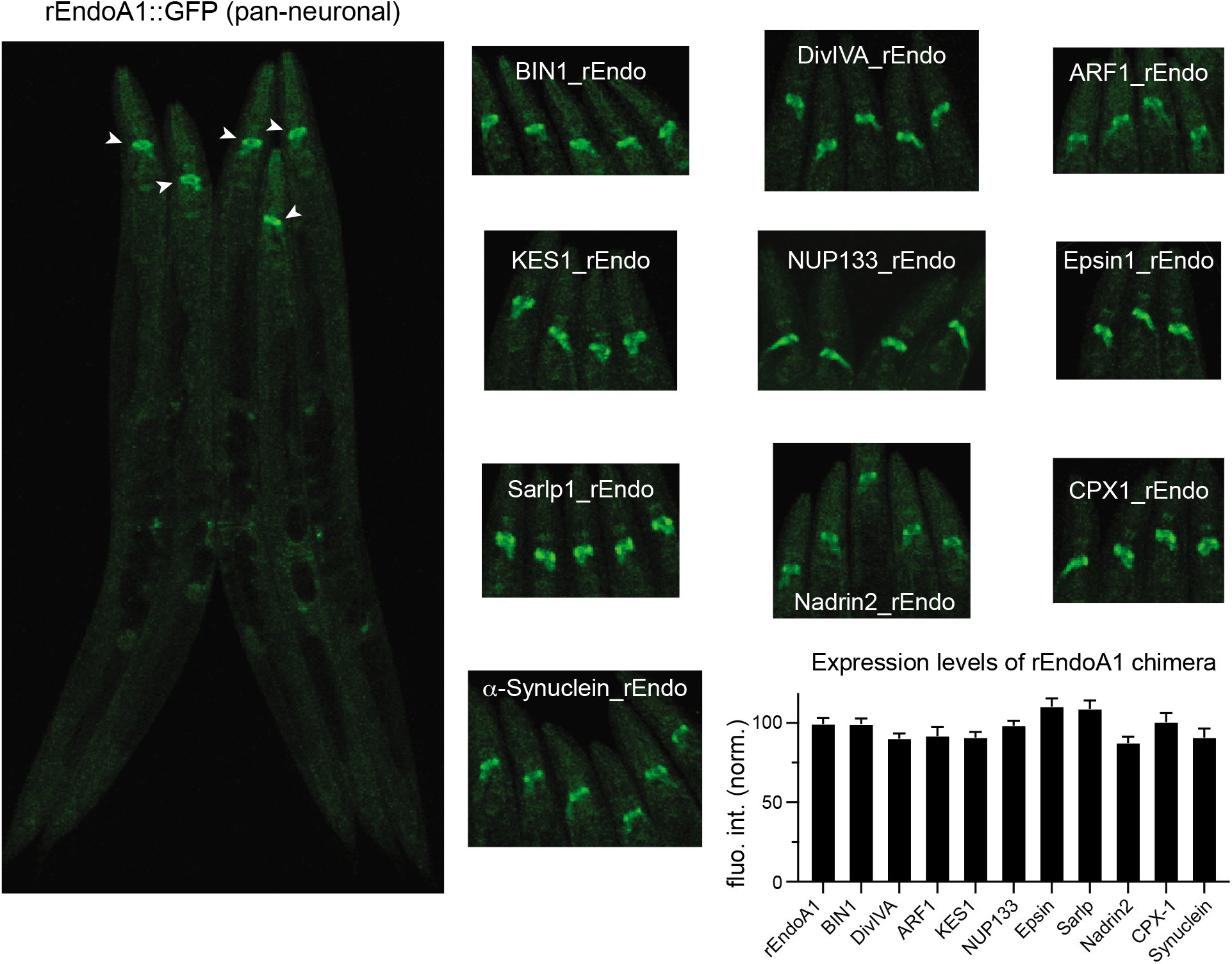

**Suppl. figure 3.**
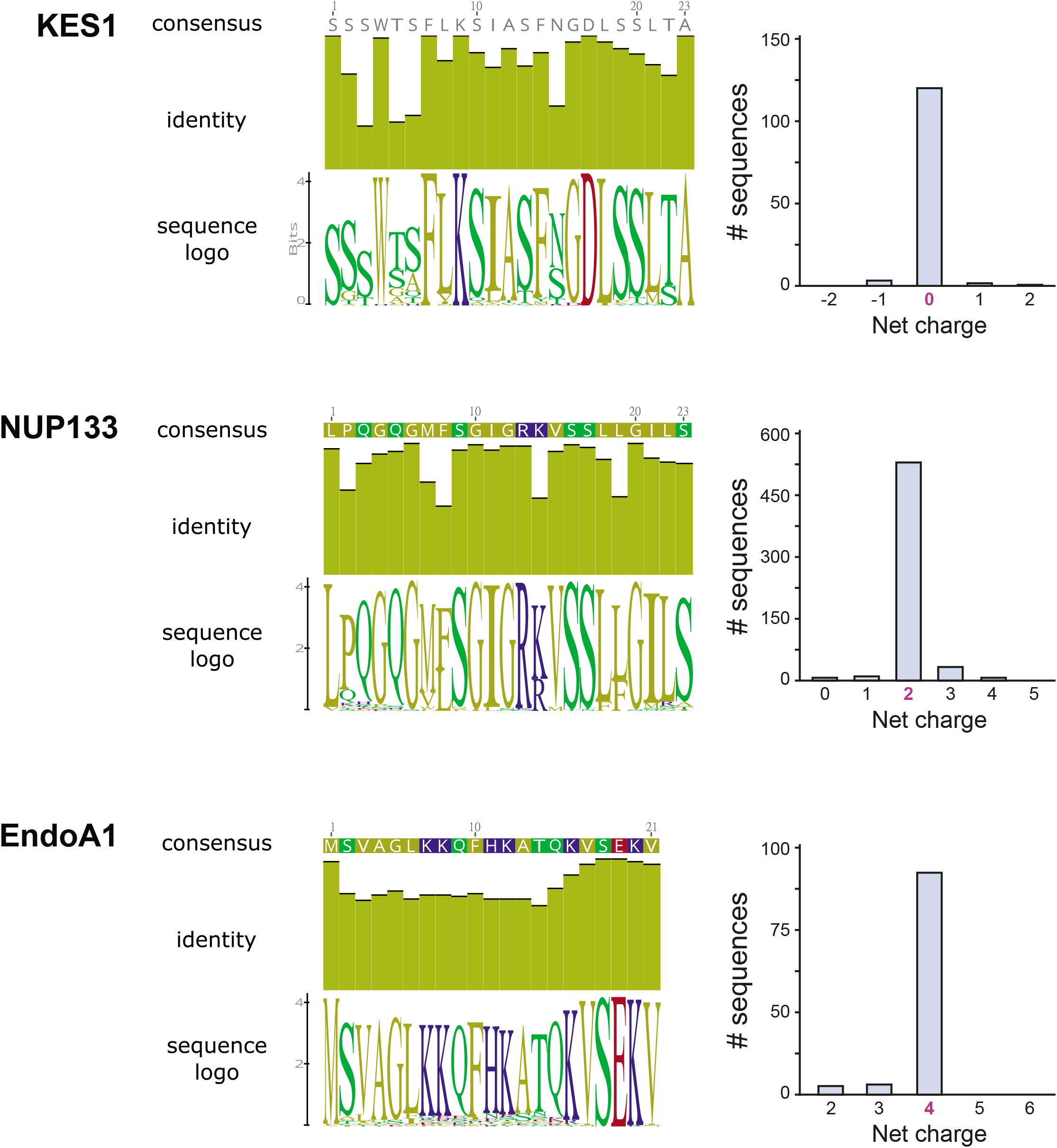

**Suppl. figure 4.**
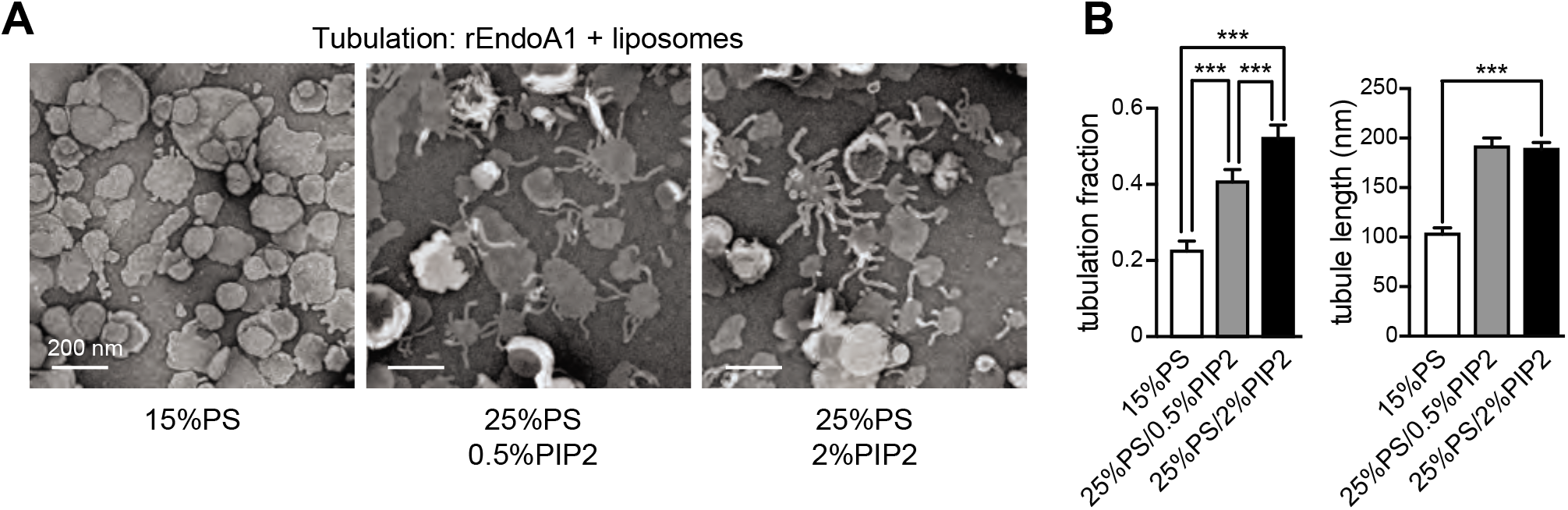

**Suppl. figure 5.**
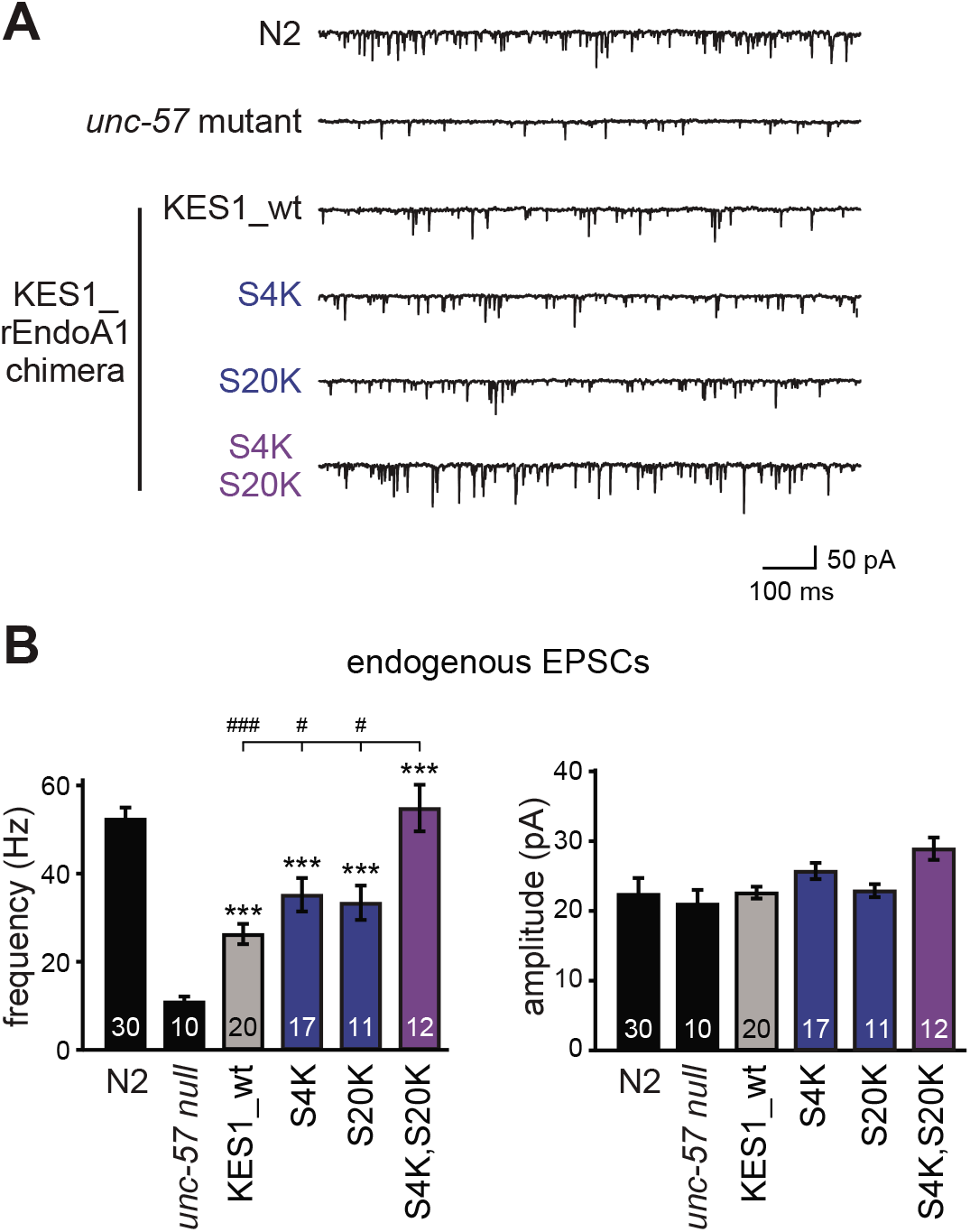

**Suppl. figure 6.**
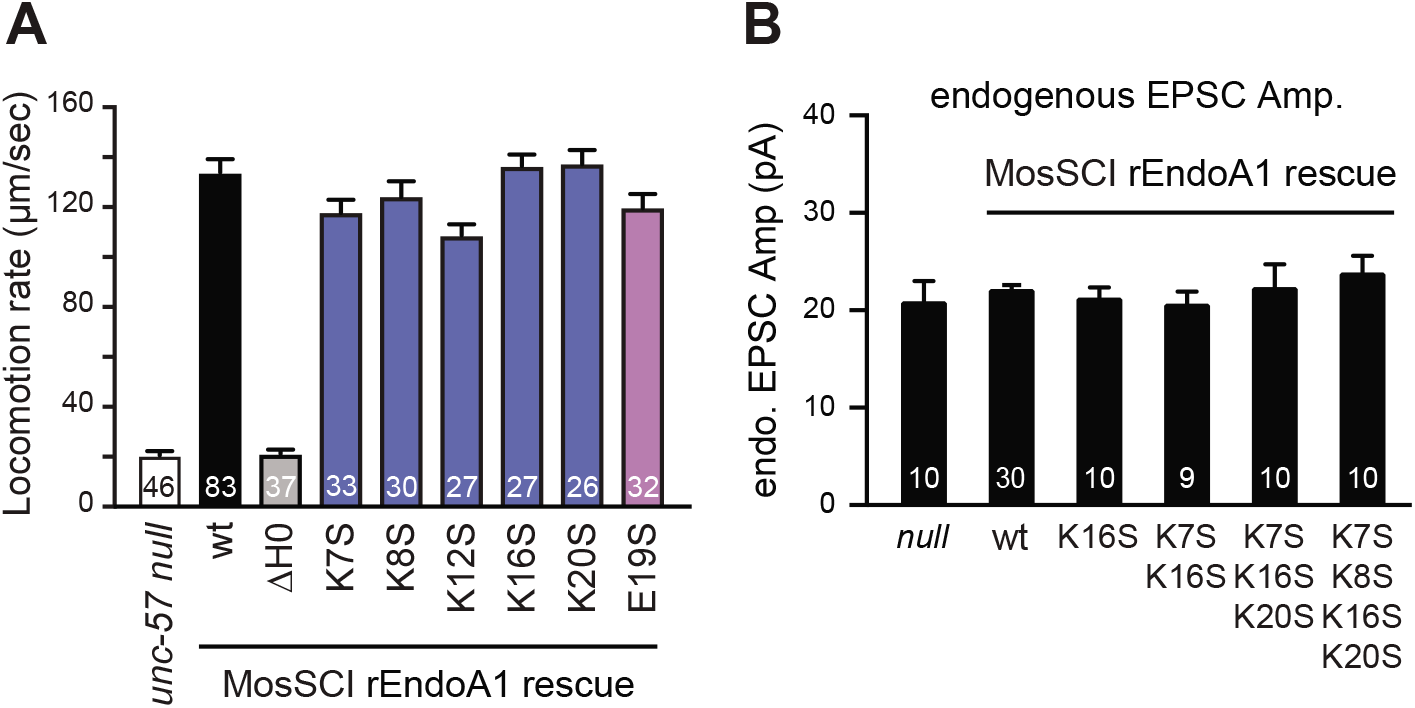

