## Supplementary material for "The endophilin curvature-sensitive motif requires electrostatic guidance to recycle synaptic vesicles *in vivo*": Suppl

**Suppl. figure 1. The function of endophilin requires its amphipathic helix H0.**

Single-copy transgenes were generated to express GFP-tagged rEndoA1 in neurons in *unc-57* mutants. ΔH0: a truncated version of rat endophilin A1 (rEndoA1) lacking amino acid residues 2-21. **(A)** Representative trajectories (15 animals) of 30s locomotion. The starting points for each trajectory were aligned (indicated by red dots) for clarity. The scale bar indicates 1 mm. **(B)** Summary data for endogenous EPSC rates and amplitudes. **(C)** Representative evoked EPSC traces (*left*) and summary data of the amplitude (*right*) were shown for indicated genotypes. ***, $p < 0.001$; n.s., not significant, when compared to *endophilin unc-57 mutants*. The number of worms analyzed for each genotype is indicated in the bar graphs. Error bars indicate SEM.

**Suppl. figure 2. GFP-tagged rEndoA1 chimeras were expressed in neurons at similar levels.**

Single-copy transgenes were generated to express GFP-tagged rEndoA1 chimeras in *C. elegans* neurons. Expression levels of rEndoA1 chimeras in neurons were estimated by measuring background-subtracted fluorescence around nerve rings (a tight axon bundle made of processes from sensory and interneurons) in *C. elegans.* White arrowheads indicated nerve rings. Fluorescence intensity of rEndoA1 chimeras was normalized to that of wild type rEndoA1. Results were plotted in the bar graph. Error bars indicate SEM.

**Suppl. figure 3. Amphipathic helices of KES1, NUP133, and Endophilin have distinct net charge properties.** The consensus sequences and the logo plots of conservation were shown in the *left* panels, for the amphipathic helices of KES1, NUP133, and Endophilin. Amino acid residues in the logo plots were colored as follows: yellow - nonpolar; green - polar; blue - basic; red - acidic. The net charge histograms of the amphipathic helices were shown in the right panels. Logo plots were generated using Geneious Prime 2021.1. Net charge was calculated for each helix using a custom script.

**Suppl. figure 4. The membrane tubulation activity of Endophilin is regulated by the abundance of anionic phospholipids. (A)** Transmission electron micrographs for liposomes (2 mM total lipids) incubated with wild type rEndoA1 (4 μM) for 5 min at room temperature. Liposomes were prepared by extrusion through filters with 0.2 μm pore size. The lipid compositions of liposomes were 15%PS/85%PC *(left)*, 25%PS/0.5%PIP2/74.5%PC *(middle)*, and 25%PS/2%PIP2/73%PC *(right)*. **(B)** Quantification of membrane tubulation. *(left)* Tubulation Fraction = (# of liposomes with extended protrusions) / (# of total liposomes). *(right)* The length of membrane tubules was determined using ImageJ. Error bars indicate SEM. ***, $p < 0.001$. One-way ANOVA followed by Dunnett's test was used for statistical analysis.

**Suppl. figure 5. S/K mutations improved the ability of KES1_rEndoA1 chimera to support endogenous synaptic activity *in vivo*. (A)** Representative traces of endogenous excitatory postsynaptic currents that were recorded at neuromuscular junctions were shown for indicated genotypes. Single-copy transgenes were introduced to drive pan-neuronal expression of KES1_rEndoA1 variants in *endophilin unc-57(e406)* mutant worms. **(B)** Summary data for endogenous EPSC rates *(left)* and amplitudes *(right)* were shown. ***, $p < 0.001$; when compared to *endophilin unc-57 mutants*. $^{###}$, $p < 0.001$; and $^{#}$, $p < 0.05$; when compared to animals rescued by S4K,S20K mutant KES1_rEndoA1. The number of worms analyzed for each genotype is indicated in the bar graphs. Error bars indicate SEM. One-way ANOVA followed by Dunnett's test was used for statistical analysis.

**Suppl. figure 6. The impact of rEndoA1 K/S mutations on locomotory and electrophysiological features. (A)** Quantification of locomotion rates. Mutant rEndoA1 chimeras with single Lys-to-Ser and Glu-to-Ser mutations were examined for their ability to restore locomotion activity in *endophilin unc-57* mutant worms. The number of worms analyzed for each genotype is shown in the bar graphs. Error bars represent SEM. **(B)** The amplitude of endogenous EPSCs was plotted for Lys-to-Ser mutants of rEndoA1. Error bars represent SEM. The number of worms analyzed for each genotype is indicated in the bar graphs.

**KEY RESOURCES TABLE**

| REAGENT OR RESOURCE | SOURCE | IDENTIFIER |
|---|---|---|
| **Bacterial** | | |
| *E. coli*: OP50 | CGC | OP50 |
| **Chemicals and Critical Commercial Assays** | | |
| PrimeSTAR GXL DNA Polymerase | TaKaRa | Cat # R050A |
| PrimeSTAR Max DNA Polymerase | TaKaRa | Cat # R045B |
| Phusion polymerase | NEB Biolabs | Cat # M0530S |
| Gateway BP Clonase II | ThermoFisher | Cat # 11789100 |
| Gateway LR Clonase II | ThermoFisher | Cat # 11791020 |
| Ni-NTA agarose beads | Qiagen | Cat. # 157030990 |
| HEPES | ThermoFisher | Cat. # BP310-500 |
| Isopropyl–d-thiogalactopyranoside (IPTG) | ThermoFisher | Cat. # BP1620-10 |
| Dithiothreitol | Boston BioProducts | Cat. # P765 |
| 2,3-butanedione monoxime (BDM) | Sigma | Cat. # B0753 |
| Agar | Apex | Cat. # 20-275 |
| QIAquick PCR Purification Kit | QIAGEN | Cat # 28104 |

| **Experimental Models: Organisms/Strains** | | |
|---|---|---|
| Wild type Bristol isolate | CGC | N2 |
| *unc-57(e406) I* | CGC | CB406 |
| *ttTi5605 II; unc-119(ed9) III* | (Frokjaer-Jensen et al., 2008) | EG4322 |
| *unc-119(ed3) III; cxTi10816 IV* | (Frokjaer-Jensen *et al.*, 2008) | EG6401 |
| *kyIs673 [Psra-6::eat-4::pHluorin]* | (Ventimiglia and Bargmann, 2017) | CX16921 |
| *unc-57(e406) I; ttTi5605 II* | This study | BJH2045 |
| *ttTi5605 II* | This study | BJH2002 |
| *cxTi10816 IV* | This study | BJH2147 |
| *unc-57(e406) I; pekSi423[snb-1p::rEndoA1::gfp::unc-54utr + neoR (+)] II* | This study | BJH2488 |
| *unc57(e406) I; pekSi435[snb-1p::rEndoA1(Gly_insertion)::gfp + hygR(+)] II* | This study | BJH2494 |
| *unc57(e406) I; pekSi436[snb-1p::rEndoA1(GlyGly_insertion)::gfp + hygR(+)] II* | This study | BJH2495 |
| *unc57(e406) I, pekSi145[snb-1p::rEndoA1(GlyGlyAla_insertion)::gfp + neoR(+)] II* | This study | BJH2127 |

| | | |
|---|---|---|
| *unc-57(e406) I; pekSi274[snb-1p::bin1_AH::rEndoA1ΔH0::gfp + neoR(+)] II* | This study | BJH3146 |
| *unc-57(e406) I; pekSi477[snb-1p::divIVA_AH::rEndoA1ΔH0::gfp + neoR(+)] II* | This study | BJH2567 |
| *unc-57(e406) I; pekSi217[snb-1p::arf1_AH::rEndoA1ΔH0::gfp + neoR(+)] II* | This study | BJH3089 |
| *unc-57(e406) I; pekSi304[snb-1p::nadrin2_AH::rEndoA1ΔH0::gfp + neoR(+)] II* | This study | BJH3176 |
| *unc-57(e406) I; pekSi181[snb-1p::kes1_AH::rEndoA1ΔH0::gfp + neoR(+)] II* | This study | BJH3054 |
| *unc-57(e406) I; pekSi210[snb-1p::nup133_AH::rEndoA1ΔH0::gfp + neoR(+)] II* | This study | BJH3082 |
| *unc-57(e406) I; pekSi273[snb-1p::epn1_AH::rEndoA1ΔH0::gfp + neoR(+)] II* | This study | BJH3145 |
| *unc-57(e406) I; pekSi479[snb-1p::sarlp_AH::rEndoA1ΔH0::gfp + neoR(+)] II* | This study | BJH2568 |
| *unc-57(e406) I; pekSi452[snb-1p::cpx-1_AH::rEndoA1ΔH0::gfp + neoR(+)] II* | This study | BJH3324 |
| *unc-57(e406) I; kyIs673[sra-6p::eat-4::pHluorin]* | This study | BJH2333 |
| *unc-57(e406) I; pekSi480[snb-1p::nup133_AH::rEndoA1ΔH0 + neoR(+)] II; kyIs673[sra-6p::eat-4::pHluorin]* | This study | BJH3352 |
| *unc-57(e406) I; pekS482[snb-1p::kes1_AH::rEndoA1ΔH0 + neoR(+)] II; kyIs673[sra-6p::eat-4::pHluorin]* | This study | BJH2569 |
| *unc-57(e406) I; pekS484[snb-1p::rEndoA1 + neoR(+)] II; kyIs673[sra-6p::eat-4::pHluorin]* | This study | BJH3356 |
| *unc-57(e406) I; pekS489[snb-1p::rEndoA1ΔH0 + neoR(+)] II; kyIs673[sra-6p::eat-4::pHluorin]* | This study | BJH3361 |
| *unc-57(e406) I; pekSi481[snb-1p::KES1_AH(S4K, S20K)::rEndoA1ΔH0 + NeoR(+)] II; kyIs673 [sra-6p::eat-4::pHluorin]* | This study | BJH3353 |

| | | |
|---|---|---|
| *unc-57(e406)I; pekSi236[snb-1p::rEndoA1(K12S)::gfp + NeoR(+)] II* | This study | BJH3108 |
| *unc-57(e406)I; pekSi262[snb-1p::rEndoA1(K16S)::gfp + NeoR(+)] II* | This study | BJH3134 |
| *unc-57(e406)I; pekSi462[snb-1p::rEndoA1(K7S,K16S)::gfp + NeoR(+)] II* | This study | BJH3344 |
| *unc-57(e406)I; pekSi473[snb-1p::rEndoA1(K7S,K16S,K20S)::gfp + NeoR(+)] II* | This study | BJH3345 |
| *unc-57(e406)I; pekSi474[snb-1p::rEndoA1(K7S,K8S,K16S,K20S)::gfp + NeoR(+)] II* | This study | BJH3346 |
| *unc-57(e406) I; pekSi387[snb-1p::kes1_AH(S2K)::rEndoA1ΔH0::gfp + neoR(+)] II* | This study | BJH2407 |
| *unc-57(e406) I; pekSi388[snb-1p::kes1_AH(S3K)::rEndoA1ΔH0::gfp + neoR(+)] II* | This study | BJH2408 |
| *unc-57(e406) I; pekSi378[snb-1p::kes1_AH(S4K)::rEndoA1ΔH0::gfp + neoR(+)] II* | This study | BJH3250 |
| *unc-57(e406) I; pekSi389[snb-1p::kes1_AH(T6K)::rEndoA1ΔH0::gfp + neoR(+)] II* | This study | BJH2409 |
| *unc-57(e406) I; pekSi390[snb-1p::kes1_AH(S7K)::rEndoA1ΔH0::gfp + neoR(+)] II* | This study | BJH2410 |
| *unc-57(e406) I; pekSi393[snb-1p::kes1_AH(S11K)::rEndoA1ΔH0::gfp + neoR(+)] II* | This study | BJH2417 |
| *unc-57(e406) I; pekSi400[snb-1p::kes1_AH(S14K)::rEndoA1ΔH0::gfp + neoR(+)] II* | This study | BJH2459 |
| *unc-57(e406) I; pekSi391[snb-1p::kes1_AH(S20K)::rEndoA1ΔH0::gfp + neoR(+)] II* | This study | BJH2411 |
| *unc-57(e406) I; pekSi392[snb-1p::kes1_AH(S21K)::rEndoA1ΔH0::gfp + neoR(+)] II* | This study | BJH2412 |
| *unc-57(e406) I; pekSi396[snb-1p::kes1_AH(S23K)::rEndoA1ΔH0::gfp + neoR(+)] II* | This study | BJH2458 |
| *unc-57(e406) I; pekSi468[snb-1p::kes1_AH(S4K,S20K)::rEndoA1ΔH0::gfp + neoR(+)] II* | This study | BJH3340 |

| | | |
|---|---|---|
| *unc-57(e406) I; pekSi477[snb-1p::divIVA_AH::rEndoA1ΔH0::gfp + neoR(+)] II* | This study | BJH2567 |
| *unc-57(e406) I; pekSi445[snb-1p::divIVA_AH(E4S,D12S,E14S)::rEndoA1ΔH0::gfp + neoR(+)] II* | This study | BJH3317 |
| *unc-57(e406) I; kyIs673 [sra-6p::eat-4::pHluorin]; pekEx225[snb-1p::divIVA _AH::rEndoA1ΔH0]* | This study | BJH1084 |
| *unc-57(e406) I; kyIs673 [sra-6p::eat-4::pHluorin]; pekEx225[snb-1p::divIVA _AH(E4S,D12S,E14S)::rEndoA1ΔH0]* | This study | BJH1085 |
| *unc-57 (e406) I; pekSi394[snb-1p::nup133_AH(K/S)::rEndoA1ΔH0::gfp + NeoR(+)] II* | This study | BJH2456 |
| *pekSi288[unc-129p::unc-57::mNeonGreen + hygR(+)] II; pekSi441[unc-129p::unc-57::mScarlet + neoR(+)] IV* | This study | BJH3313 |
| *pekSi288[unc-129p::unc-57::mNeonGreen + hygR(+)] II; pekSi431[unc-129p::rEndoA1_H0::unc-57ΔH0::mScarlet + neoR(+)] IV* | This study | BJH3303 |
| *pekSi288[unc-129p::unc-57::mNeonGreen + hygR(+)] II; pekSi432[unc-129p::kes1_AH::unc-57ΔH0::mScarlet + neoR(+)] IV* | This study | BJH3304 |
| *pekSi288[unc-129p::unc-57::mNeonGreen + hygR(+)] II; pekSi443[unc-129p::nup133_AH::unc-57ΔH0::mScarlet + neoR(+)] IV* | This study | BJH3315 |
| **Recombinant DNA** | | |
| Plasmid: pPD49.26 vector | | |
| Plasmid: pCFJ356 | | |
| Plasmid: pCFJ350 | | |

| | | |
|---|---|---|
| Plasmid: pET28a | | |
| Plasmid: *snb-1p::rEndoA1::gfp::unc-54utr* | This study | BJP-A734 |
| Plasmid: *snb-1p::rEndoA1(Gly_insertion)::gfp::unc-54utr* | This study | BJP-C734 |
| Plasmid: *snb-1p::rEndoA1(GlyGly_insertion)::gfp::unc-54utr* | This study | BJP-C735 |
| Plasmid: *snb-1p::rEndoA1(GlyGlyAla_insertion)::gfp::unc-54utr* | This study | BJP-A501 |
| Plasmid: *snb-1p::bin1_AH::rEndoA1ΔH0::gfp::unc-54utr* | This study | BJP-A729 |
| Plasmid: *snb-1p::divIVA_AH::rEndoA1ΔH0::gfp::unc-54utr* | This study | BJP-C828 |
| Plasmid: *snb-1p::arf1_AH::rEndoA1ΔH0::gfp::unc-54utr* | This study | BJP-A730 |
| Plasmid: *snb-1p::kes1_AH::rEndoA1ΔH0::gfp::unc-54utr* | This study | BJP-A725 |
| Plasmid: *snb-1p::nup133_AH::rEndoA1ΔH0::gfp::unc-54utr* | This study | BJP-A726 |
| Plasmid: *snb-1p::epn1_AH::rEndoA1ΔH0::gfp::unc-54utr* | This study | BJP-A727 |
| Plasmid: *snb-1p::sarlp_AH::rEndoA1ΔH0::gfp::unc-54utr* | This study | BJP-C827 |
| Plasmid: *snb-1p::rEndoA1::unc-54utr* | This study | BJP-C845 |
| Plasmid: *snb-1p::kes1_AH::rEndoA1ΔH0::unc-54utr* | This study | BJP-C846 |
| Plasmid: *snb-1p::nup133_AH::rEndoA1ΔH0::unc-54utr* | This study | BJP-C847 |
| Plasmid: *snb-1p::rEndoA1ΔH0::unc-54utr* | This study | BJP-C915 |
| Plasmid: *snb-1p::kes1_AH(S4K,S20K)::rEndoA1ΔH0::unc-54utr* | This study | BJP-C848 |

| | | |
|---|---|---|
| Plasmid: *snb-1p::rEndoA1(K7S,K16S)::gfp::unc-54utr* | This study | BJP-C783 |
| Plasmid: *snb-1p::rEndoA1(K7S,K16S,K20S)::gfp::unc-54utr* | This study | BJP-C782 |
| Plasmid: *snb-1p::rEndoA1(K7S,K8S,K16S,K20S)::gfp::unc-54utr* | This study | BJP-C781 |
| Plasmid: *snb-1p::rEndoA1_(K7S)::unc-54utr* | This study | BJP-C952 |
| Plasmid: *snb-1p::rEndoA1_(K8S)::unc-54utr* | This study | BJP-C953 |
| Plasmid: *snb-1p::rEndoA1_(K20S)::unc-54utr* | This study | BJP-C954 |
| Plasmid: *snb-1p::kes1_AH(S2K)::rEndoA1ΔH0::gfp::unc-54utr* | This study | BJP-C584 |
| Plasmid: *snb-1p::kes1_AH(S3K)::rEndoA1ΔH0::gfp::unc-54utr* | This study | BJP-C585 |
| Plasmid: *snb-1p::kes1_AH(T6K)::rEndoA1ΔH0::gfp::unc-54utr* | This study | BJP-C586 |
| Plasmid: *snb-1p::kes1_AH(S7K)::rEndoA1ΔH0::gfp::unc-54utr* | This study | BJP-C587 |
| Plasmid: *snb-1p::kes1_AH(S11K)::rEndoA1ΔH0::gfp::unc-54utr* | This study | BJP-C515 |
| Plasmid: *snb-1p::kes1_AH(S14K)::rEndoA1ΔH0::gfp::unc-54utr* | This study | BJP-C588 |
| Plasmid: *snb-1p::kes1_AH(S20K)::rEndoA1ΔH0::gfp::unc-54utr* | This study | BJP-C589 |
| Plasmid: *snb-1p::kes1_AH(S21K)::rEndoA1ΔH0::gfp::unc-54utr* | This study | BJP-C590 |
| Plasmid: *snb-1p::kes1_AH(S23K)::rEndoA1ΔH0::gfp::unc-54utr* | This study | BJP-C591 |
| Plasmid: *snb-1p::kes1_AH(S4K,S20K)::rEndoA1ΔH0::gfp::unc-54utr* | This study | BJP-C772 |
| Plasmid: *snb-1p::divIVA_AH::rEndoA1ΔH0::unc-54utr* | This study | BJP-C929 |

| | | |
|---|---|---|
| Plasmid: *snb-1p::divIVA_AH(E4S,D12S,E14S)::rEndoA1ΔH0::unc-54utr* | This study | BJP-C930 |
| Plasmid: *snb-1p::nup133_AH(K/S)::rEndoA1ΔH0::gfp::unc-54utr* | This study | BJP-C518 |
| Plasmid: *his6::sumo::rEndoA1* | This study | BJP-C702 |
| Plasmid: *his6::sumo::rEndoA1ΔH0* | This study | BJP-C703 |
| Plasmid: *his6::sumo::nup133_AH::rEndoA1ΔH0* | This study | BJP-A761 |
| Plasmid: *his6::sumo::kes1_AH::rEndoA1ΔH0* | This study | BJP-A763 |
| Plasmid: *his6::sumo::kes1_AH(S4K,S20K)::rEndoA1ΔH0* | This study | BJP-C951 |
| **Software and imagining equipment** | | |
| Excel | Microsoft | N/A |
| Prism 9 | GraphPad Prism | N/A |
| WormLab Imaging System | MBF Bioscience, VT, USA | N/A |
| Olympus FV-1000 Confocal Microscope | Olympus Company | N/A |
| SnapGene 5.0 | SnapGene | N/A |
| Igor Pro | Wavemetrics, OR, USA | N/A |

| | |
|---|---|
| MATLAB | MathWorks, Inc., Natick, MA |

Note: rEndoA1 indicates rat endophilin A1

### LEAD CONTACT AND MATERIALS AVAILABILITY

### DATA AND CODE AVAILABILITY

The datasets generated during, and/or analyzed during the current study are available from the corresponding author upon reasonable request.

### EXPERIMENTAL MODEL AND SUBJECT DETAILS

Worm strains were maintained at 22°C on NGM (nematode growth medium) agar plates seeded with Escherichia coli OP50 bacteria according to standard protocols (Brenner 1974). Unless otherwise specified, all experiments were carried out using day 1 adult hermaphrodites. Mutant and transgenic alleles were backcrossed at least 4 times against N2 Bristol strains.

### METHOD DETAILS

#### Molecular Biology

DNA plasmids were constructed using the Multisite Gateway system (Invitrogen, Waltham, MA, USA) and Gibson assembly protocols (Gibson et al., 2009). All constructs were sequence verified. cDNA encoding rat endophilinA1 (rEndoA1) was amplified as previously described (Bai *et al.*, 2010). The promoters used in this study are *Psnb-1* (3kb, for rescue experiments) and *Punc-129* (2.6kb, for imaging analyses). These promoters were cloned into modified pCFJ150 and pCFJ356 vectors

(Frokjaer-Jensen *et al.*, 2008) along with the cDNA of gene of interest. To build chimeric endophilin variants, the H0 helix (residues 2-21, SVAGLKKQFHKATQKVSEKV) of rat endophilin A1 was replaced with known amphipathic helices (listed below). DNA constructs encoding chimeric endophilin variants were generated using overlap extension PCR strategy (Higuchi et al., 1988). To construct plasmids for recombinant protein expression, DNA fragments encoding genes of interest were inserted into a modified PET28A vector (Novagen, Madison, WA) containing a His6 tag to facilitate purification.

BIN1_AH [8-33, *H. sapiens*]

*VTAGKIASNVQKKLTRAQEKVLQKL*

*5’-gtgactgctggaaaaatagcttcaaatgtccagaagaagttgactcgtgcccaagagaaggttctccaaaaactc-3’*

ARF1_AH [1-17, *H. sapiens*]

MGNIFANLFKGLFGKKE

*5’-atgggtaacatctttgccaacctgtttaaaggtctttttggcaagaaggag-3’*

NUP133_AH [245-267, *H. sapiens*]

LPQGQGMLSGIGRKVSSLFGILS

*5’-cttccacaaggtcagggcatgctcagtggaattggtagaaaagtatcttcattgttcggaatcctttca-3’*

KES1_AH [7-29, *S. cerevisiae*]

SSSWTSFLKSIASFNGDLSSLSA

5’-tcatcatcctggacatctttcctgaaatctatcgcttcctttaatggtgatttgtcgtcgctgagtgcc-3’

Sarlp_AH [1-23]

MAGWDIFGWFRDVLASLGLWNKH

5’-*atggccggatgggacatcttcggatggttccgtgacgtccttgcctcccttggactttggaacaagcac*-3’

EPN1_AH [3-16, *H. sapiens*]

TSSLRRQMKNIVHN

5'-*acgtcttcccttcgaagacagatgaaaaacatagtacataac*-3'

CPX-1_AH [115-135, *C. elegans*]

LGLTEQVEKAKTMATGAFETV

5'-*ctcgggctcacggagcaagtcgaaaaagcgaaaactatggcgaccggcgccttcgagaccgtc*-3'

divIVA_AH [25-41, *B. subtilis*]

VNEFLAQVRKDYEIVLR

5'-*gtcaacgagttccttgcccaagtccgtaaggactacgagatcgtcctccgt*-3'

**Recombinant Protein Production**

Recombinant proteins were expressed as N-terminal his6-SUMO-tagged fusion proteins in the BL21(DE3) *E. coli* strain. Protein purification and tag removal were carried out using published protocols (Poudel *et al.*, 2016). Briefly, the bacteria culture was grown in Luria Broth medium at 37°C. When $OD_{600}$ reached 1.0, isopropyl-D-thiogalactopyranoside was added to the bacterial culture (0.2mM) to induce protein expression. The bacterial culture was further grown overnight at 15°C. Bacteria harvested by centrifugation was lysed by a microfluidizer in the lysis buffer (20 mM HEPES, pH 8.0, 300 mM NaCl, 15 mM imidazole). Proteins were purified using NiNTA-agarose beads (Qiagen, Valencia, CA), and then eluted with a lysis buffer plus 250 mM imidazole. Recombinant endophilinA1 variants were expressed as his6-SUMO-tagged fusion proteins. The SUMO protease ULP1 (ubiquitin-like-specific protease 1) was used to cleave off his6-SUMO tags, resulting in endophilin A1 variants with native N-terminal amino acid compositions. His6-SUMO fragments cleaved off by ULP1 were removed using NiNTA-agarose beads (Qiagen, Valencia, CA). Purified proteins were dialyzed against the HEPES buffer (50 mM HEPES, pH 7.4, 150mM NaCl) plus 1 mM DTT. Proteins were stored at 4°C.

**Transgenes and Germline Transformation**

Transgenic strains were generated by microinjection. Mos1-mediated single-copy transgene insertion methods were used to produce animals carrying single-copy transgenes (Frokjaer-Jensen et al., 2012; Frokjaer-Jensen *et al.*, 2008). Mos1 target sites used in this study are *ttTi4391* (chromosome I) and *cxTi10816* (chromosome IV). Transgenic worms were outcrossed at least four times.

**Worm Tracking and Locomotion Analysis**

Young adult animals (day 1) were picked to 10 cm agar plates with no bacterial lawn (20 worms per plate). Imaging began 1 hour after worms were transferred. Worm crawling on the agar surface was recorded for 30 seconds using the WormLab Imaging System (MBF Bioscience, VT, USA). The center of mass was determined for each animal using custom object-tracking software developed using ImageJ (Schneider et al., 2012). Average speed was determined for each animal.

**Confocal Microscopy and Analysis**

Animals were immobilized with 2,3-Butanedione monoxamine (30 mg/ml; Sigma-Aldrich), and were mounted on 2% agarose pads for imaging. Fluorescence images were collected on an inverted Olympus IX81 microscope, using a laser scanning confocal imaging system (Olympus FluoView FV1000) and an Olympus UPLSAPO 60x 1.4 NA Oil immersion objective (at 5x zoom). GFP and mNeonGreen were excited using a 488 nm Argon laser and mScarlet was excited using a 559 nm diode pumped solid state laser. Rat endophilinA1 (rEndoA1) variants used in the rescue experiments were C-terminally tagged with GFP. Expression levels of rEndoA1 variants were estimated by measuring background-subtracted fluorescence around the *C. elegans* nerve ring. Dorsal nerve cords were imaged to determine synaptic and axonal distribution of rEndoA1 variants. Fluorescence images were analyzed using custom software in IGOR Pro (WaveMetrics, Lake Oswego, OR) (Burbea et al.,

2002; Dittman and Kaplan, 2006). Images of fluorescent slides (Chroma Technology Group, VT) were collected daily to monitor the laser stability, and the dorsal cord fluorescence was normalized to the slide fluorescence value. Statistical significance was determined using one-way ANOVA or Student's t-test and all values reported are means ± SEM.

**VGLUT-pHluorin Imaging and Analysis**

Worms carrying the *kyIs673* [*sra-6p::eat-4::pHluorin*] transgene were used for imaging VGLUT-pHluorin fluorescence as previously described (Ventimiglia and Bargmann, 2017). Animals were loaded into custom-built PDMS microfluidic chambers designed to deliver stimuli under a fluorescent microscope (Chronis et al., 2007). To prevent movement, animals were paralyzed with 1mM tetramisole hydrochloride. NaCl stimulus (500mM) was prepared fresh daily in S. Basal Buffer. The ASH axons were imaged on an inverted Leica DMi8 microscope through a Leica PL APO 63x 1.40 NA oil immersion objective, onto an Andor iXon Life 888 EMCCD camera, using Leica LAS-X software. Animals were allowed to acclimate for 5 minutes in microfluidic chambers before imaging, and were adapted to blue light for 90 seconds. No more than 3 trials were performed per animal. The time duration for all recording trials was 100 seconds, and the stimulus was presented for 10 seconds, starting at 30 seconds after imaging began. Images were captured at 5 frames per second.

Imaging analysis was performed as previously described (Ventimiglia and Bargmann, 2017). Briefly, images were corrected for x-y drift, and images with substantial z-drift were discarded. The axon was divided into 3x3 pixel ROIs by hand, and background ROIs to the left and right of each axon ROI were selected. Intensity and pixel measurements of the axon and background were extracted from those ROIs. Background and bleaching artifacts were corrected. VGLUT-pHluorin has slow bleaching kinetics, whereas the background exhibits significant bleaching. To correct for this, background bleaching was linearly modeled through a ‘polyfit’ function in MATLAB and the linear fit was subtracted from the background signal. After bleaching correction, background correction was performed by subtracting fluctuations in the background signal from the fluorescence traces. Basal fluorescence of VGLUT-pHluorin was obtained by averaging the first 10 seconds of each ROI for each trial after background correction. Decay curves were fitted and ΔF/F was determined as previously described. All imaging experiments for a given condition or observation were repeated on at least two separate days using independently prepared buffers and stimuli. The number of trials per animal and the total number of animals are reported along with the total trial number in the figure legends.

**Liposome Preparation**

Lipids were stored at −20°C. POPC (1-palmitoyl-2-oleoyl-sn-glycero-3-phosphocholine, designated as PC) was purchased from Larodan Inc, DOPS (1,2-di-oleoyl-sn-glycero-3-phospho-l-serine, designated as PS) was obtained from NOF America Corporation, and PI(4,5)P2 (brain L-α-phosphatidylinositol-4,5-bisphosphate, designated as PIP2) was purchased from Avanti Polar Lipids. Lipids dissolved in chloroform were mixed in a glass tube. The lipid mixture was dried under a stream of nitrogen gas for 30 min, and residual solvent was removed under vacuum (Labconco Free Zone 2.5 lyophilizer) for 2 hours. The dried lipid film was then rehydrated in the HEPES buffer (50 mM HEPES, 150 mM NaCl, pH 7.4), followed by a brief sonication to homogenize the mixture. The resulting lipid suspension was then extruded 21 times through 200 nm pore-size polycarbonate membranes, using a Mini extruder (Avanti Polar lipids) to form unilamellar vesicles.

**Transmission Electron Microscope for Visualizing Synaptic Vesicle**

Approximately 10 adult hermaphrodites were quickly loaded into a 100 μm freezing chamber containing space-filling bacteria. These worms were frozen instantaneously at ~ -180 °C in a Leica EM PACT2 system. The frozen worms were fixed in a Leica EM AFS2 machine using 1% osmium in 0.1% UA in acetone fixative, then embedded in Eponate 12 from Ted Pella, Inc. Serial sections were cut at a thickness of 40 nm using an ultracut E microtome, collected on pioloform-covered slotted grids (notchnum 1x2 mm oval) from Ted Pella, Inc., and counterstained in 6% aqueous uranyl acetate for 1.5 hrs, followed by Reynolds lead citrate for 7 min. Images were obtained on a Talos L120C TEM transmission electron microscope (Thermo Fisher Scientific Inc, USA) operating at 120KV.

Micrographs were collected using the Ceta-M 4k x 4k high-resolution camera (Thermo Fisher Scientific Inc, USA). Synapse profiles were used to count the number of synaptic vesicles. Each profile represents a single section that passes through the dense projection. P values were generated using one-way ANOVA followed by Dunnett's test.

**Electrophysiology**

Day-1 young adult hermaphrodite *C. elegans* were dissected as previously described (Dong et al., 2015; Richmond et al., 1999). Animals were immobilized on Sylgard-coated coverslips using tissue adhesive glue (Histoacryl Blue, Braun), and were dissected in extracellular solution via a dorsolateral incision using a sharpened tungsten needle. After removing gonad and intestines by suction through a glass pipette, the cuticle flap was turned and gently glued down to expose the underlying ventral nerve cord and body-wall-muscle quadrants. The worm prep was then mounted onto a fixed-stage upright microscope (BX51WI, Olympus) equipped with a 60x water-immersion objective lens. The integrity of the anterior ventral body wall muscle and the ventral nerve cord were visually examined via the DIC microscopy, and ventral muscle cells were patched using fire-polished 2-5 MΩ resistant borosilicate pipettes (World Precision Instruments, USA). Whole-cell patch clamp recordings were carried out at 20°C. Body-wall muscle cells were voltage clamped at -60 mV to record postsynaptic currents. The currents were recorded in the whole-cell configuration from muscle cells using an amplifier (EPC-10; HEKA, Germany). Extracellular solution contained (in mM) 150 NaCl, 5 KCl, 1 CaCl2, 5 MgCl2, 10 glucose and 10 HEPES, titrated to pH 7.3 with NaOH, 330 mOsm with sucrose. Internal solution contained 135 CH3O3SCs, 5 CsCl, 5 MgCl2, 5 EGTA, 0.25 CaCl2, 10 HEPES and 5 Na2ATP, adjusted to pH 7.2 using CsOH. Evoked EPSC responses were induced by applying a 0.4 ms, 30 μA pulse, generated by a stimulus isolator (A365, WPI), through a borosilicate pipette (~2 MΩ) placed in close apposition to the ventral nerve cord. Series resistance was compensated to 70% for the evoked EPSC recording. All chemicals were purchased from Sigma. Data were sampled at 10 kHz using Patchmaster (HEKA), following low-pass filtering at 2 kHz. The number of animals for each experiment was specified in Figure legends.

### Alignment of Protein Sequences

For each amphipathic helix-containing protein (EndoA1 – *R. norvegicus*, NP_446387; KES1 – *S. cerevisiae*, NP_015180; Nup133 – *H. sapiens*, NP_060700), the helix sequence, plus 50 amino acids on either end, where possible (some helices were terminal), was used as a BLASTP query to identify orthologs within the NR database. These longer protein sequences were aligned using MAFFT (PMID: 12136088), and the aligned helix sequences were extracted. For each set of helix sequences, sequences annotated as the gene of interest were extracted, and splice isoforms and duplicate sequences were removed to generate an alignment with a single sequence from each sampled species. Logo plots were generated using Geneious Prime 2021.1. Net charge was calculated for each helix using a custom script.

### Negative-Stain Electron Microscopy

Sample preparation for negative stain was carried out as previously described with modifications (Farsad *et al.*, 2001; Poudel *et al.*, 2016). Briefly, liposomes (total lipids 2 mM, 0.2 μm in diameter) were incubated with endophilin variants (4 μM) in the HEPES buffer (50mM HEPES, 150 mM NaCl, pH7.4) for 5 min at room temperature. Nickel grids were glow-discharged for 40 seconds, after which grids were incubated with samples for 2 minutes. Samples were fixed using 0.5x Karnovsky's fix. Grids were then washed with 1 drop of 0.1 M cacodylate buffer, followed with a 4 drop $H_2O$ wash. A drop of 1% uranyl acetate was touched on the sample. The grids were then carefully dragged through dry filter papers and put in a desiccator overnight to dry. Images were collected using JEOL TEM 1400 and Talos L120C Transmission Electron Microscopes. All measurements were made using ImageJ. Ten images were selected for analysis by following random number tables generated by Google Sheets.

### Statistics

Student's t-test was used to compute significance for a single pairwise comparison. One-way ANOVA followed by Tukey or Dunnett's test was used for multiple comparisons. $p < 0.05$ was considered to be statistically significant (*$p < 0.05$, **$p < 0.01$, ***$p < 0.001$). Statistical analysis and graphing were carried out using Prism8 (GraphPad), Igor Pro (WaveMetrics), Clampfit (Molecular Devices), FluoView (Olympus), and Excel (Microsoft). Data were presented as the mean ± standard error (SEM).
